# Genotype imputation improves GWAS resolution and facilitates candidate gene identification in pearl millet

**DOI:** 10.64898/2026.10.16.759592

**Authors:** Kota Kambara, Shashi Kumar Gupta, Daisuke Tsugama

## Abstract

Pearl millet (*Cenchrus americanus*; syn. *Pennisetum glaucum* (L.) R. Br.) is an important cereal crop mainly grown in semi-arid areas. However, in pearl millet, genes underlying many agronomic traits remain unknown. Our previous genome-wide association study (GWAS) was based primarily on genotype data derived from restriction site-associated DNA sequencing (RAD-seq), which contained a high proportion of missing genotypes and may have limited mapping resolution. To address this limitation, we used whole-genome sequencing (WGS)-derived genotype data from 21 elite lines as a reference panel to impute missing genotypes in the RAD-seq-derived dataset. Genotype imputation reduced the proportion of missing genotypes from 82.42% to 0% and increased the mean density of non-missing genotypes from 569.61 to 1,158.09 genotypes/Mb. Leave-one-out validation yielded a mean genotype concordance of 89.9 ± 4.6%. GWAS using the imputed dataset showed more localized association signals for several traits. Homologs of rice *SHORT INTERNODES1*, Arabidopsis *FLOWERING LOCUS Y*, and rice *EPIDERMAL PATTERNING FACTOR 2* were identified as candidates for genes regulating culm height, days to panicle emergence, and awn growth, respectively. These results demonstrate that reference-panel-based genotype imputation can increase the utility of reduced-representation genotype datasets and facilitate candidate gene identification in pearl millet.

**Key message:** Imputation of missing genotypes in RAD-seq-derived genotype data using a WGS-derived reference panel improved GWAS resolution and facilitated candidate gene identification for agronomic traits in pearl millet.

## Introduction

Pearl millet (*Cenchrus americanus*; syn. *Pennisetum glaucum* (L.) R. Br.) is an important cereal crop mainly cultivated in semi-arid regions such as sub-Saharan Africa and India. Its tolerance to drought, high temperature, and low-nutrient conditions makes it particularly important in environments where water and agricultural inputs are limited. Genomic resources for pearl millet have expanded substantially since the release of its reference genome and the subsequent development of an improved genome assembly (Varshney et al. 2017; Ramu et al. 2023). These resources have enabled genome-wide association studies (GWAS) to identify genomic regions associated with yield-related traits, rhizosheath formation, and other agronomic traits (de la Fuente Cantó et al. 2022; Rouamba et al. 2023). However, the genetic basis of many agronomic traits in pearl millet remains poorly understood.

B– and R-lines developed by the International Crops Research Institute for the Semi-Arid Tropics (ICRISAT) are major parental materials used in pearl millet breeding (Mula et al. 2007). In our previous study, 16 traits were evaluated in these elite lines, and GWAS was performed using genotypes derived primarily from restriction site-associated DNA sequencing (RAD-seq; Varshney et al. 2017; Kambara et al. 2024). The GWAS identified a chromosome 3 region associated with pigmentation of the shoot basal part and a homolog of *PHENYLALANINE AMMONIA-LYASE 2* (*PAL2*) as a candidate gene. For culm height, strongly associated polymorphisms were detected across a broad region of chromosome 4, making it difficult to identify a specific candidate gene underlying the trait. For the remaining traits, no specific genomic region showed a strong association (Kambara et al. 2024). Thus, although this population showed phenotypic variation for multiple agronomic traits, genes underlying most of these traits, including culm height, days to panicle emergence, and awn growth, remained unidentified.

Reduced-representation sequencing approaches such as RAD-seq provide cost-effective genome-wide genotype information for large populations, but the resulting datasets can contain substantial proportions of missing genotypes because of limited sequencing depth, variation in restriction sites, and uneven representation of genomic regions (Poland and Rife 2012). Missing genotypes and uneven marker coverage can reduce the power and resolution of GWAS. Genotype imputation can infer unobserved genotypes using observed marker and haplotype information from the study population or a densely genotyped reference panel (Marchini et al. 2007; Chan et al. 2016). It can thereby increase effective marker density and improve the power and resolution of subsequent association analyses. However, no reference panel for genotype imputation has been established in pearl millet.

Here, we show that imputing these missing genotypes using a densely genotyped reference panel improves GWAS resolution and thereby facilitates the identification of candidate genes underlying agronomic traits. For this, we constructed a reference panel using whole-genome sequencing (WGS) data from 21 elite pearl millet lines and used it to impute missing genotypes in the RAD-seq-derived dataset. These reference panel and GWAS data can be resources to promote pearl millet genomics and breeding.

## Materials and methods

### Plant materials

Seeds of the pearl millet lines listed in Supplementary Table S1 were provided by the International Crops Research Institute for the Semi-Arid Tropics (ICRISAT, India). Seeds were surface-sterilized with 70% (v/v) ethanol for 1 min, followed by treatment with a sterilization solution containing 5% sodium hypochlorite and 0.05% (v/v) Tween-20 for 30 min, and were then sown on medium containing 1.5% (w/v) agar. These seeds were incubated in a growth chamber at 30 °C under a 12-h light/12-h dark photoperiod for 1 week and subsequently transplanted into pots containing soil supplemented with fertilizer. Leaves were collected from 3-week-old plants and stored at −80 °C until DNA extraction.

### DNA extraction, whole-genome sequencing, and genotyping

Genomic DNA was extracted from leaves of 3-week-old plants using the ISOSPIN Plant DNA kit (Nippon Gene Co., Ltd., Tokyo, Japan). The extracted genomic DNA was sent to Macrogen Japan Corp. (Tokyo, Japan) for WGS. Sequencing was performed on an Illumina NovaSeq X Plus platform to generate 150-bp paired-end reads, yielding approximately 50 Gb of sequence data per sample. The sequenced reads were mapped to the Tift23D_2_B_1_-P1-P5 (Tift) pearl millet reference genome assembly (Ramu et al. 2023) using Bowtie2 v2.5.1 (Langmead and Salzberg 2012), and sequence variants were called using minipileup v1.4b (https://github.com/lh3/minipileup) to obtain genotypic data. RAD-seq data for 81 elite lines used in Kambara et al. (2024), excluding the 21 lines subjected to WGS in this study, were obtained from the National Center for Biotechnology Information (NCBI) Sequence Read Archive (SRA) (Varshney et al. 2017; accession numbers are listed in Supplementary Table S1). These RAD-seq reads were mapped to the pearl millet reference genome described above using Bowtie2, and sequence variants were called using minipileup.

### Field trial

A field trial was conducted at the Institute for Sustainable Agro-ecosystem Services, The University of Tokyo, Japan (35.74°N, 139.54°E), following the procedures used for the 2022 field experiment described by Kambara et al. (2024). A total of 91 pearl millet lines were grown from July 2023 to January 2024 (Supplementary Table S2). Days to panicle emergence were recorded as the number of days from sowing to panicle emergence.

### Genotype imputation and GWAS

Genotype data obtained from the WGS data of the 21 pearl millet lines were used as a reference panel for genotype imputation. Variant sites containing missing genotype calls were excluded from the reference panel using VCFtools v0.1.17 (Danecek et al. 2011). Missing genotypes in the RAD-seq-derived genotype dataset were imputed using Beagle v5.4 (Browning et al. 2018), with the reference panel. The imputed RAD-seq genotype data were then combined with the WGS-derived genotype data and polymorphisms with minor allele frequency (MAF) < 0.05 were removed. The resulting genotype dataset contained 102 lines, comprising the 21 reference-panel lines and 81 additional lines. In parallel, the original RAD-seq genotype dataset without genotype imputation was combined with the WGS-derived genotype data to generate the non-imputed genotype dataset. Genome-wide association analyses were performed using the gaston package v1.6 (Dandine-Roulland and Perdry 2018) in R with phenotypic data from the 2023 field trial described above or from Kambara et al. (2024). Associations were tested using a linear mixed model (LMM), with genetic relatedness among lines accounted for using a genomic relationship matrix calculated from standardized genome-wide genotype data. The genome-wide significance thresholds were determined using the method of Li and Ji (2005), implemented in the poolr package v1.2 in R (Cinar and Viechtbauer 2022), and were set at *P* < 4.68e-5 for the non-imputed genotype dataset and *P* < 4.66e-5 for the imputed genotype dataset. Manhattan plots were generated using the qqman package v0.1.9 (Turner 2018) in R. For each major GWAS peak, genes located within ±1 Mb of the lead single-nucleotide polymorphism (SNP) were extracted, and their functional annotations based on Arabidopsis and rice orthologs were obtained from TGIF-DB (Tsugama and Takano 2021). To characterize sequence variation in candidate regions in greater detail, including indels and complex short variants, variants were independently called from the WGS data of the 21 lines using Octopus v0.7.4 (Cooke et al. 2021). Candidate variants located within these genes were extracted, and their genotypes among the 21 lines were compared with the corresponding phenotypic values. Phenotypic values were compared among genotype or diplotype groups using the Tukey–Kramer test for culm height and days to panicle emergence, and Dunn’s test with Holm correction for awn growth. Pairwise linkage disequilibrium (LD) among SNPs in the regions surrounding major association peaks was calculated and visualized as *r*^2^ using LDBlockShow (Dong et al. 2021).

### Leave-one-out validation of genotype imputation accuracy

Imputation accuracy was assessed by leave-one-out validation using the 20 reference-panel lines for which both WGS and RAD-seq data were available (Supplementary Table S3). In each iteration, one focal line was removed from the reference panel, and its WGS-derived genotypes were retained as the validation truth set. The RAD-seq-derived genotypes of the focal line were then imputed using the remaining 20 reference lines. Genotype concordance was calculated as the proportion of evaluated sites at which the imputed diploid genotype exactly matched the corresponding WGS-derived genotype.

### Quantitative reverse transcription-PCR (qRT-PCR)

Stem tissues of 3-week-old pearl millet plants were frozen and ground to a fine powder using a mortar and pestle. Total RNA was extracted from the ground tissues using a NucleoSpin RNA Plant kit (Macherey-Nagel, Düren, Germany). cDNA was synthesized from 1 μg of total RNA using PrimeScript™ IV 1st strand cDNA Synthesis Mix (Takara Bio, Shiga, Japan). Quantitative PCR was performed using a StepOne Real-Time PCR System (Thermo Fisher Scientific, Waltham, MA, USA) and TB Green Premix Ex Taq II (Tli RNaseH Plus) (Takara Bio). *PgSHI1* transcript levels were compared between B843 and R102560, whereas *PgFY* transcript levels were compared between B94333 and B00222. Three independent biological replicates were analyzed for each line. Relative expression levels were calculated using the comparative cycle threshold (Ct) method, with *PgEF1*α as the reference gene. Primers for *PgEF1*α were designed according to Shivhare and Lata (2016), and all primer sequences used for qRT-PCR are listed in Supplementary Table S4. Relative expression levels were compared between lines using Welch’s *t*-test, with *P* < 0.05 considered statistically significant.

### Cloning of *PgSHI1*

The full-length coding sequences (CDSs) of *PgSHI1* were amplified by PCR from cDNA prepared from stem tissues of B843 and R102560 as described above, using KOD FX Neo (TOYOBO, Osaka, Japan). The primers were designed based on the CDS of *PgSHI1* (*dpca4g296570.840*) annotated in the Tift reference genome (Ramu et al. 2023) and are listed in Supplementary Table S5. The PCR products were separated by agarose gel electrophoresis and purified using a FastGene Gel/PCR Extraction Kit (Nippon Genetics, Tokyo, Japan). The purified PCR products were cloned into the pBS-35SMCS-GFP vector (Tsugama et al. 2012) digested with SpeI (New England Biolabs, Ipswich, MA, USA) using In-Fusion Snap Assembly Master Mix (Takara Bio, Shiga, Japan), yielding pBS-35S-PgSHI1(B843)-GFP and pBS-35S-PgSHI1(R102560)-GFP.

### Subcellular localization analysis of PgSHI1

pBS-35S-PgSHI1(B843)-GFP and pBS-35S-PgSHI1(R102560)-GFP were introduced into onion epidermal cells together with pBS-35SMCS-mCherry (Tsugama et al. 2013) using a particle bombardment device (Tsugama and Takano 2020; Tsugama 2026). After bombardment, the cells were incubated at room temperature for 24 h, and GFP and mCherry fluorescence signals were observed as described previously (Tsugama and Takano 2020).

### Examination of gene expression using the Grass Expression Atlas

Expression profiles for *dpca4g296570.840* (*PgSHI1*), *dpca6g389000.840* (*PgFY*), and *dpca3g255860.840* (*PgEPF2*) were examined across the available pearl millet tissue and developmental-stage datasets using the Grass Expression Atlas (GExA; Kambara et al. 2026).

### Phylogenetic analysis

Protein sequences of Arabidopsis and rice were obtained from The Arabidopsis Information Resource (TAIR; Berardini et al. 2015) and Rice Genome Annotation Project (RGAP), release 7 (Kawahara et al. 2013), respectively. Protein sequences of *Setaria italica*, *Sorghum bicolor*, *Zea mays*, and *Hordeum vulgare* were obtained from Phytozome (Goodstein et al. 2012). Candidate homologs were identified by BLASTP (Camacho et al. 2009) searches with an E-value cutoff of 1e-20. For PgFY (dpca6g389000.840) and PgSHI1 (dpca4g296570.840), amino acid sequences were analyzed using NGPhylogeny.fr (Lemoine et al. 2019), with MAFFT (Katoh and Standley 2013) for sequence alignment, BMGE (Criscuolo and Gribaldo 2010) for alignment trimming, and PhyML (Guindon et al. 2010) for maximum-likelihood phylogenetic inference with 1,000 bootstrap replicates. For PgEPF2 (dpca3g255860.840), amino acid sequences were aligned using MAFFT, trimmed using trimAl (Capella-Gutiérrez et al. 2009), and subjected to maximum-likelihood phylogenetic analysis using IQ-TREE (Nguyen et al. 2015), with the best-fit substitution model selected by ModelFinder (Kalyaanamoorthy et al. 2017) and branch support assessed using 1,000 bootstrap replicates.

### Prediction of RNA secondary structures, protein structures, and signal peptides

RNA secondary structures of *PgFY* and *PgSHI1* were predicted using MXfold2 (Sato et al. 2021). For the B00222– and B94333-type *PgFY* alleles, predicted 500-nt sequences extending from 50 nt upstream to 450 nt downstream of the start codon were used as input sequences. For the B843– and R102560-type *PgSHI1* alleles, the predicted CDSs were used as input sequences. Putative pearl millet homologs of IPA1 and FCA were identified by BLASTP searches against the predicted pearl millet protein sequences using the amino acid sequences of rice OsIPA1 (LOC_Os08g39890) and OsFCA (LOC_Os09g03610), respectively, as queries. The proteins encoded by *dpca4g302490.840* and *dpca7g511740.840*, which showed the highest sequence similarity to OsIPA1 and OsFCA, respectively, were designated PgIPA1 and PgFCA. Three-dimensional structures of the B843– and R102560-type PgSHI1 proteins and the B94333– and B00222-type PgFY proteins were predicted using AlphaFold 3 (Abramson et al. 2024). The structures of PgSHI1–PgIPA1 and PgFY–PgFCA complexes containing each PgSHI1 or PgFY type were also predicted using AlphaFold 3. Protein–protein interface area, solvation free-energy gain upon interface formation (ΔiG), and the numbers of intermolecular hydrogen bonds and salt bridges in the predicted protein complex structures were obtained from the PDB-ePISA analysis (Krissinel and Henrick 2007). The signal peptide and its cleavage site in PgEPF2 were predicted using the SignalP 6.0 web server (Teufel et al. 2022).

## Results

### Genotype imputation improves genotype completeness and GWAS resolution

Of the 102 lines included in this study, 21 lines were selected as the reference panel for genotype imputation to represent the phenotypic diversity of the population based on principal component analysis (PCA) (Supplementary Fig. S1 and Supplementary Table S1). Missing genotypes in the remaining RAD-seq-derived genotype data were then imputed using the 21-line reference panel. After imputation, the proportion of missing genotypes decreased from 82.42% to 0% (Table 1). Although the total number of retained markers decreased from 6,024,440 to 2,153,178, the mean density of non-missing genotypes per line increased from 569.61 to 1,158.09 genotypes/Mb (Table 1). Thus, genotype imputation substantially increased genotype completeness.

**Table 1.** Summary of genotype datasets before and after imputation used for genome-wide association analyses.

|  | Before imputation | After imputation |
| --- | --- | --- |
| No. of markers | 6024440 | 2153178 |
| Marker density (markers/Mb)<sup>a</sup> | 3240.25 | 1158.09 |
| Mean density of non-missing genotypes per line (genotypes/Mb)<sup>b</sup> | 569.61 | 1158.09 |
| Missing genotype rate (%) | 82.42 | 0 |

<sup>a</sup>Marker density was calculated by dividing the total number of SNP markers by the combined length of chromosomes 1–7.
<sup>b</sup>Mean density of non-missing genotypes per line was calculated by dividing the number of non-missing genotypes in each line by the combined length of chromosomes 1–7 and then averaging across the 102 lines.

The accuracy of the genotype imputation was assessed by leave-one-out validation. For 20 of the 21 reference-panel lines, RAD-seq data were publicly available (Supplementary Table S1), allowing direct comparison between WGS-derived and imputed genotypes. In each validation, one of these lines was excluded from the reference panel, and its RAD-seq-derived genotypes were imputed using the remaining 20 lines. On average, 3.96 million genotypes per line were evaluated. Genotype concordance ranged from 76.6% to 95.7%, with a mean ± SD of 89.9 ± 4.6% (Supplementary Table S3 and Supplementary Fig. S2). Overall, the imputed genotypes showed high concordance with the WGS-derived genotypes. We next examined genetic relationships among the 102 lines and performed GWAS using the imputed genotype dataset. PCA of the imputed genotype dataset revealed no clear genetic clustering among lines used for GWAS (Supplementary Fig. S3). GWAS for the 13 traits previously analyzed in Kambara et al. (2024) detected significant association signals for multiple traits (Figs. 1–4 and Supplementary Fig. S4). Several association signals in GWAS appeared narrower after imputation than before, particularly for days to panicle emergence, culm height, and panicle length (Fig. 1). Notably, the broad association signal for culm height detected on chromosome 4 using the RAD-seq-derived genotype dataset became a narrower and more distinct peak after imputation (Fig. 1). For pigmentation of the shoot basal part, a significant association signal was detected on chromosome 3, overlapping the region previously reported by Kambara et al. (2024). Significant association signals were also detected for awn growth, pigmentation of nodes, leaf curliness, panicle width, lodging, vigor at the vegetative stage, culm thickness, milligrams per seed, and plant height at 45 days after sowing, with signals occurring at distinct genomic regions across multiple chromosomes (Figs. 1–4 and Supplementary Fig. S4). Candidate genes were subsequently investigated for culm height, days to panicle emergence, and awn growth (Figs. 2–4).

**Fig. 1.**
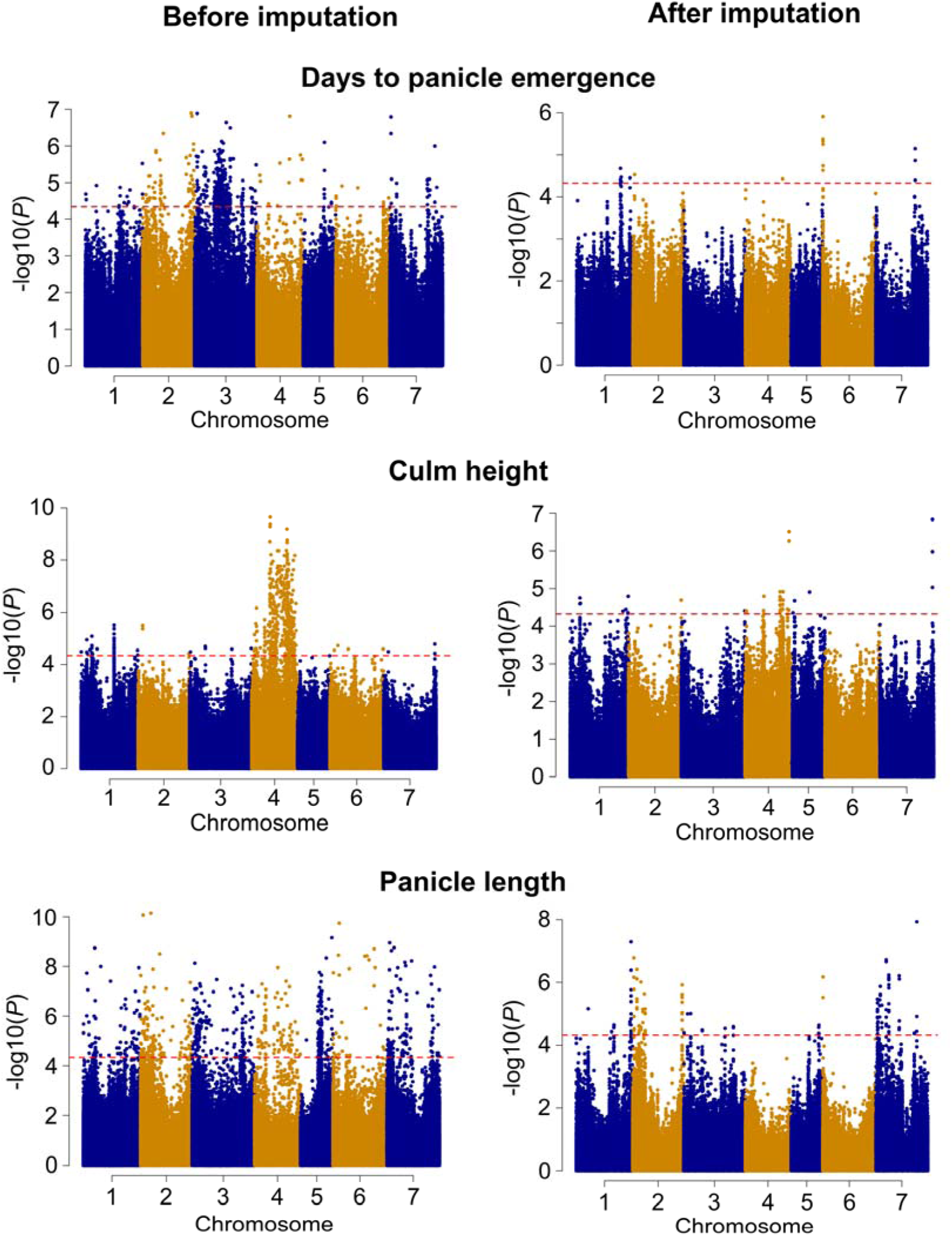
GWAS results before and after genotype imputation. Manhattan plots for days to panicle emergence, culm height, and panicle length obtained using the genotype datasets before imputation (left) and after imputation (right). The x-axis represents chromosome position, and the y-axis represents−log10(*P*). Red dashed lines indicate the genome-wide significance thresholds calculated based on the method of Li and Ji (2005), corresponding to *P* < 4.68e-5 before imputation and *P* < 4.66e-5 after imputation

**Fig. 2.**
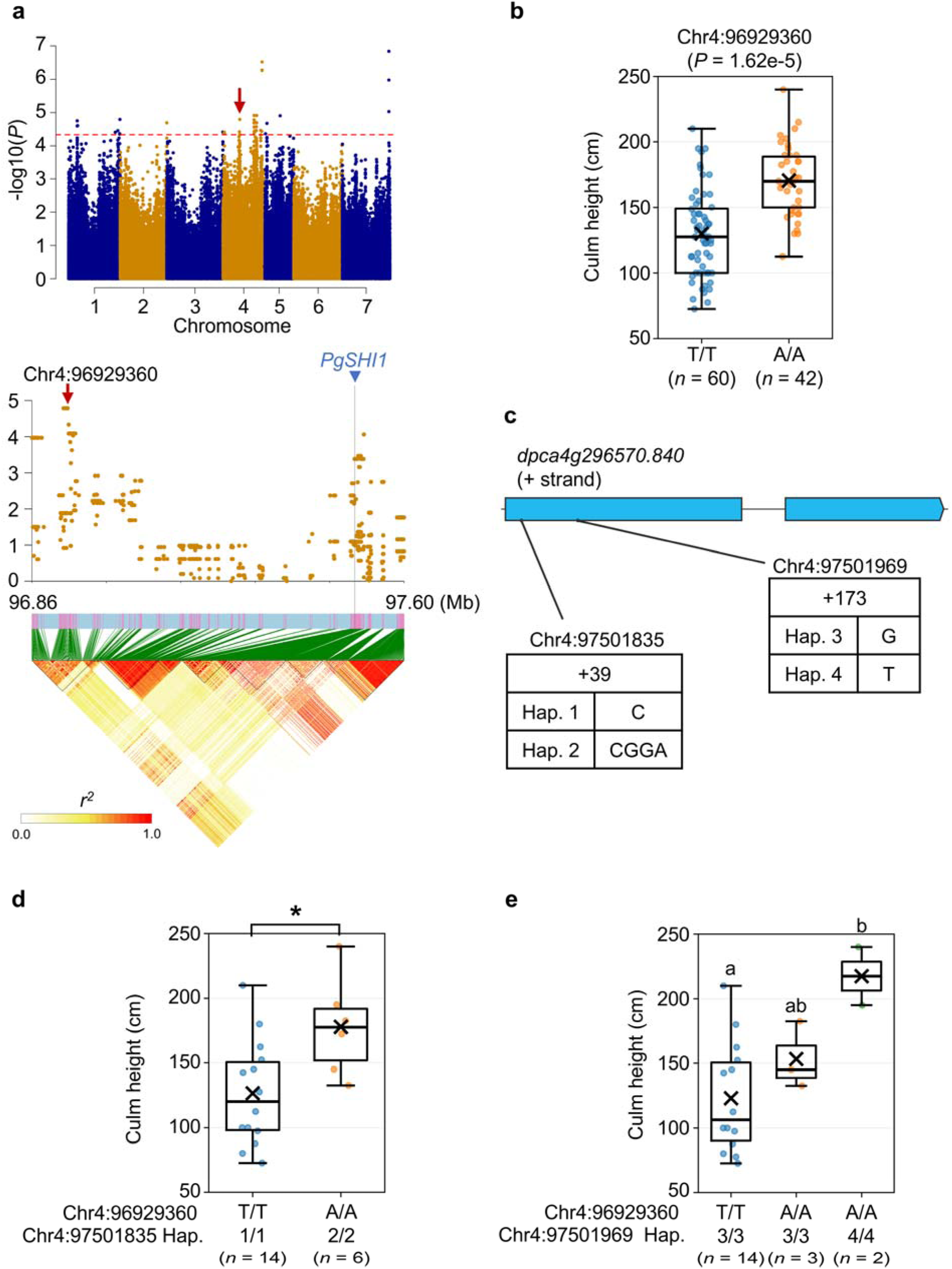
Identification of *PgSHI1* as a candidate for a gene regulating culm height in pearl millet. **a** A Manhattan plot for culm height (upper panel) and a regional association plot with linkage disequilibrium (LD) around the chromosome 4 association peak (lower panel). The red dashed line indicates the genome-wide significance threshold (*P* < 4.66e-5). The red arrow indicates the lead SNP at Chr4:96,929,360, and the blue triangle indicates the position of *PgSHI1* (*dpca4g296570.840*). Pairwise LD is shown as *r*². **b** Culm height of the 102 lines classified by genotype at the lead SNP Chr4:96,929,360. The phenotype and genotype data are those used in the GWAS shown in **a**. **c** Schematic representation of *PgSHI1* and the coding variants at Chr4:97,501,835 and Chr4:97,501,969. Boxes indicate exons, with untranslated regions (UTRs) shown in white and coding regions shown in blue; black lines indicate introns. Hap. 2 contains a 3-bp GGA insertion at Chr4:97,501,835, and Hap. 4 contains the G58V missense variant at Chr4:97,501,969. **d** Culm height of the 20 lines classified according to genotype at the GWAS lead SNP Chr4:96,929,360 and diplotype at Chr4:97,501,835. Diplotypes at Chr4:97,501,835 were determined by reanalysis of the WGS data. Genotype calls at Chr4:97,501,835 were unavailable for B09444. The asterisk indicates a significant difference (*P* < 0.05; Tukey–Kramer test). **e** Culm height of the 19 lines classified by genotype at the GWAS lead SNP Chr4:96,929,360 and diplotype at Chr4:97,501,969. Diplotypes at Chr4:97,501,969 were determined by reanalysis of the WGS data. Genotype calls at Chr4:97,501,969 were unavailable for B09333 and B94333. Individual points represent individual lines; crosses indicate means, horizontal lines indicate medians, and boxes indicate interquartile ranges. Sample sizes (*n*) are indicated below each group. Different lowercase letters indicate significant differences among groups (*P* < 0.05; Tukey–Kramer test)

**Fig. 3.**
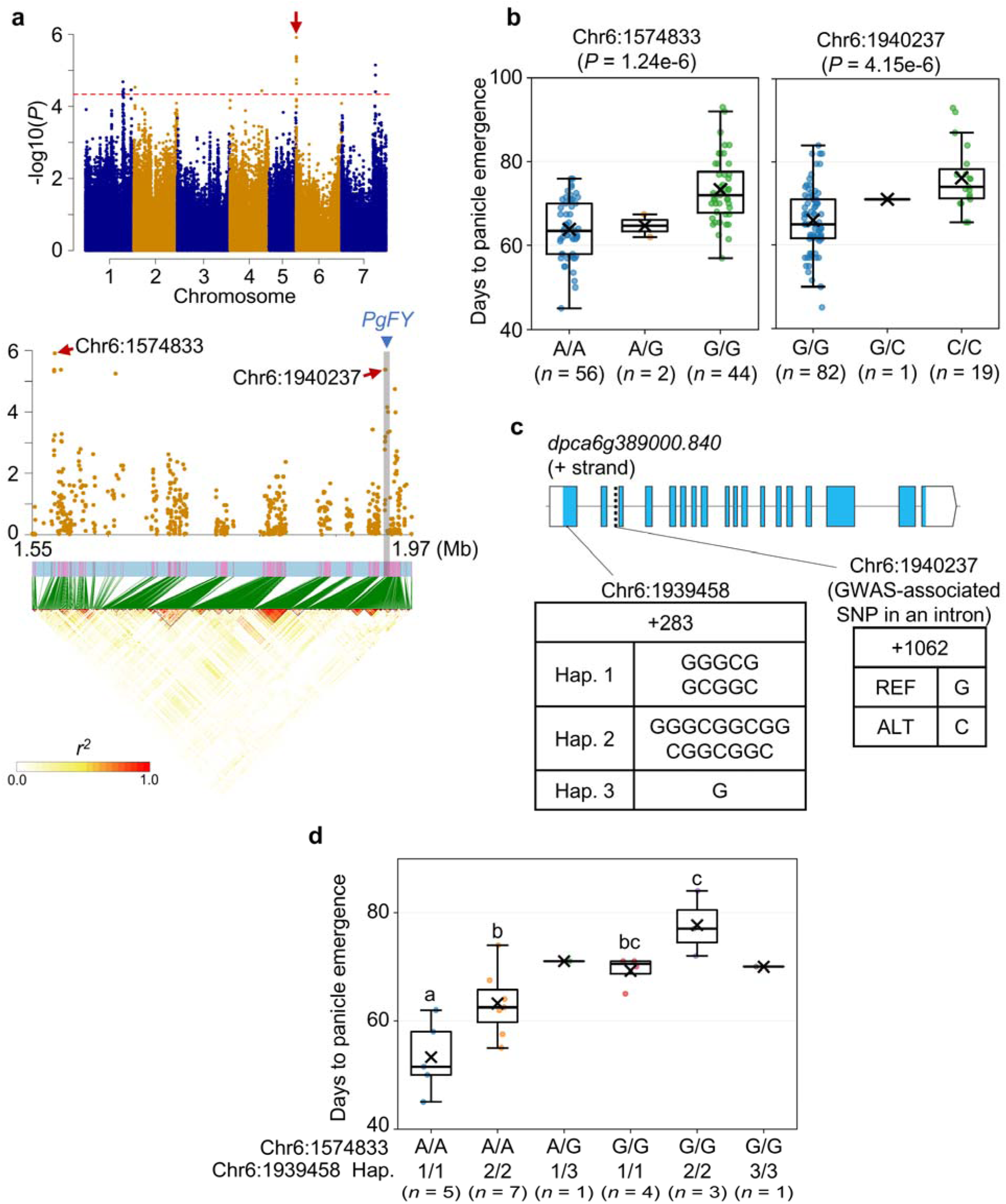
Identification of *PgFY* as a candidate for a gene regulating days to panicle emergence in pearl millet. **a** A Manhattan plot for days to panicle emergence (upper panel) and a regional association plot with linkage disequilibrium (LD) around the chromosome 6 association peak (lower panel). The red dashed line indicates the genome-wide significance threshold (*P* < 4.66e-5). Red arrows indicate the associated SNPs at Chr6:1,574,833 and Chr6:1,940,237, and the blue triangle indicates the position of *PgFY* (*dpca6g389000.840*). Pairwise LD is shown as *r*². **b** Days to panicle emergence of the 102 lines classified by genotype at Chr6:1,574,833 (left) or Chr6:1,940,237 (right). The phenotype and genotype data are those used in the GWAS shown in **a**. **c** Schematic representation of *PgFY* and sequence variants at Chr6:1,939,458 and Chr6:1,940,237. Boxes indicate exons, with untranslated regions (UTRs) shown in white and coding regions shown in blue; black lines indicate introns. At Chr6:1,939,458, Hap. 2 carries a 6-bp insertion and Hap. 3 carries a 9-bp deletion relative to Hap. 1. Chr6:1,940,237 is a GWAS-associated SNP located in an intron of *PgFY*. **d** Days to panicle emergence of the 21 lines classified by genotype at the GWAS lead SNP Chr6:1,574,833 and diplotype at Chr6:1,939,458. Diplotypes at Chr6:1,939,458 were determined by reanalysis of the WGS data from the 21 lines. Individual points represent individual lines; crosses indicate means, horizontal lines indicate medians, and boxes indicate interquartile ranges. Sample sizes (*n*) are indicated below each group. Different lowercase letters indicate significant differences among groups (*P* < 0.05; Tukey–Kramer test). Groups represented by a single line were excluded from statistical comparisons

**Fig. 4.**
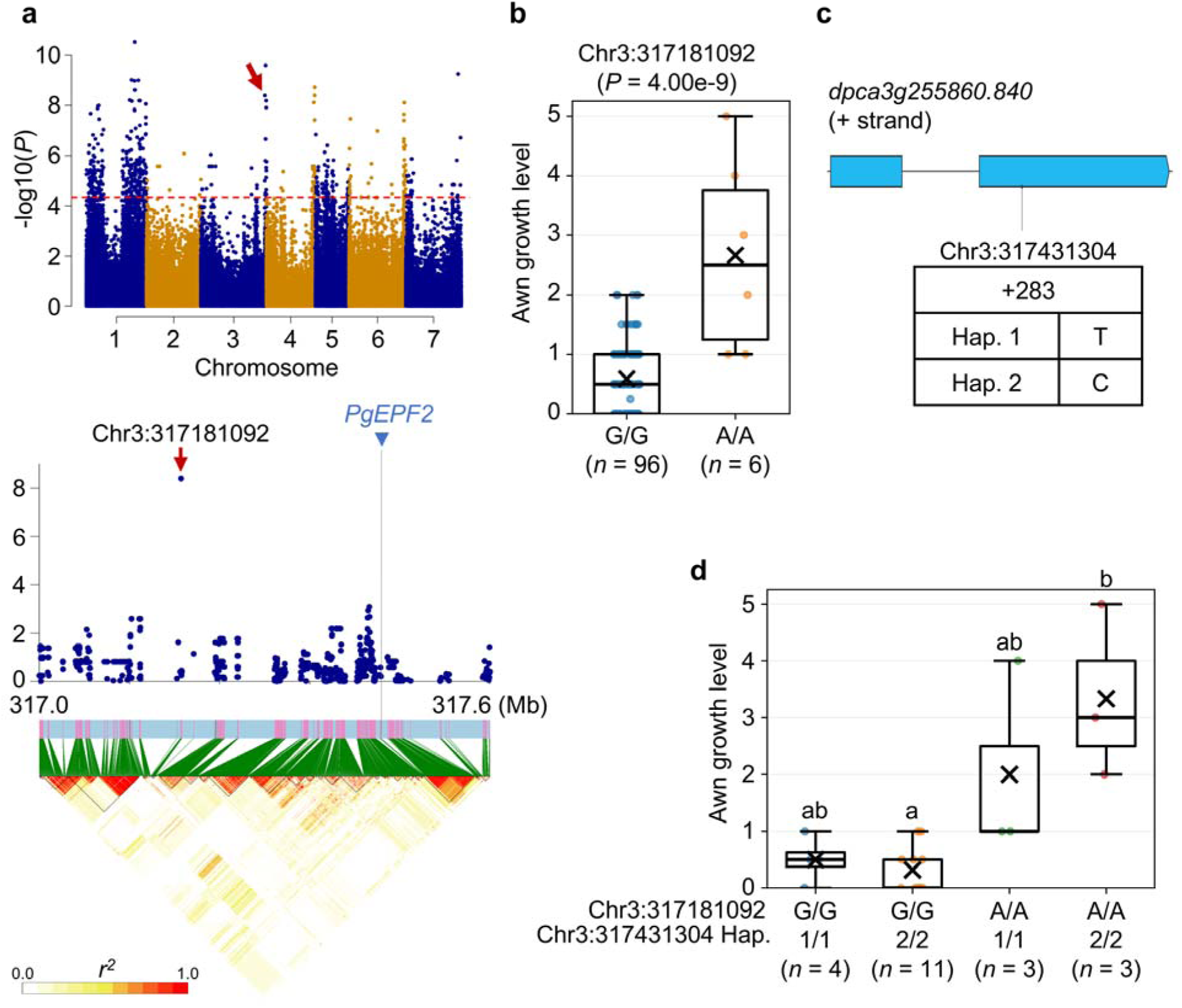
Identification of *PgEPF2* as a candidate for a gene regulating awn growth in pearl millet. **a** A Manhattan plot for awn growth (upper panel) and a regional association plot with linkage disequilibrium (LD) around the chromosome 3 association peak (lower panel). The red dashed line indicates the genome-wide significance threshold (*P* < 4.66e-5). The red arrow indicates the lead SNP at Chr3:317,181,092, and the blue triangle indicates the position of *PgEPF2* (*dpca3g255860.840*). Pairwise LD is shown as *r*². **b** Awn growth levels of the 102 lines, classified by genotype at the lead SNP Chr3:317,181,092. The phenotype and genotype data are those used in the GWAS shown in **a**. **c** Schematic representation of *PgEPF2* and the missense variant at Chr3:317,431,304 causing the M57T substitution. Boxes indicate exons, with untranslated regions (UTRs) shown in white and coding regions shown in blue; black lines indicate introns. Hap. 1 and Hap. 2 represent the two alleles at this site. **d** Awn growth levels of the 21 lines classified by genotype at the GWAS lead SNP Chr3:317,181,092 and *PgEPF2* CDS diplotype at Chr3:317,431,304. Diplotypes at Chr3:317,431,304 were determined by reanalysis of the WGS data from the 21 lines. Individual points represent individual lines; crosses indicate means, horizontal lines indicate medians, and boxes indicate interquartile ranges. Sample sizes (*n*) are indicated below each group. Different lowercase letters indicate significant differences among groups (*P* < 0.05; Dunn’s test with Holm correction)

### *PgSHI1* is a candidate for a gene regulating culm height in pearl millet

For culm height, an associated polymorphism was detected on chromosome 4 at 96,929,360 (Chr4:96,929,360; *P* = 1.62e-5) (Fig. 2a and 2b). In the proximity of the associated polymorphism, *dpca4g296570.840* was identified, which encodes a protein with high sequence similarity to rice SHORT INTERNODES1 (OsSHI1; LOC_Os09g36160). OsSHI1 is a member of the SHORT INTERNODES (SHI) transcription factor family and is involved in the regulation of plant architecture and plant height in rice (Duan et al. 2019; Duan et al. 2023). Phylogenetic analysis showed that dpca4g296570.840 clustered with OsSHI1 within the SHI family (Supplementary Fig. S5). In the GExA expression database (Kambara et al. 2026), *dpca4g296570.840* transcripts were found to be abundant in samples from florets, stalks, tillering tissues, and stems (Supplementary Fig. S6). Expression of *OsSHI1* in young panicles and axillary buds has also been reported (Duan et al. 2019). Based on these analyses, *dpca4g296570.840* was designated *PgSHI1* and considered a candidate for a gene regulating culm height in pearl millet.

Variants in the CDS of *PgSHI1* were next investigated. None of the variants in the CDS included in the GWAS marker set was significantly associated with culm height. Because the marker set excluded variants with missing genotype calls among the 21 lines for a reference panel and did not capture longer indels, we re-examined the WGS data from these 21 lines to identify CDS variants. Three variants were identified within the *PgSHI1* CDS: a 3-bp in-frame insertion of GGA at Chr4:97,501,835 (Hap. 2), a missense variant at Chr4:97,501,969 causing a Gly58Val substitution (G58V; Hap. 4), and a missense variant at Chr4:97,502,491 causing a Val198Met substitution (V198M; Hap. 6) (Fig. 2c and Supplementary Fig. S7). To examine the relationship between these variants and culm height, the 21 lines were classified according to their genotypes at the GWAS lead SNP (Chr4:96,929,360) and at each of the three variant sites in the CDS. At Chr4:97,501,835, lines with the Hap. 2/Hap. 2 diplotype had greater culm height than those with Hap. 1/Hap. 1 (Fig. 2d). At Chr4:97,501,969, lines with the A/A genotype at the GWAS lead SNP and the Hap. 4/Hap. 4 diplotype had significantly greater culm height than those with the T/T genotype and the Hap. 3/Hap. 3 diplotype, whereas culm height did not differ significantly between the two diplotypes among lines with the A/A genotype at the lead SNP (Fig. 2e). At Chr4:97,502,491, the V198M substitution was detected in two of the 21 lines, with one heterozygous and one homozygous line. Because each V198M-containing diplotype was represented by only one line, its association with culm height could not be statistically evaluated (Supplementary Fig. S7). Among the 20 lines with genotype calls at Chr4:97,501,835, the 3-bp insertion was completely concordant with the A/A genotype at the GWAS lead SNP. Together, these results suggest that the 3-bp insertion in *PgSHI1* is associated with culm height.

We next examined whether these three variants were associated with differences in *PgSHI1* expression or PgSHI1 protein properties. R102560 was homozygous for the 3-bp insertion and G58V and also carried V198M, whereas B843 carried none of the three variants (Supplementary Figs. S7 and S8). The expression level of *PgSHI1* in the stem region did not differ significantly between R102560 and B843 (Supplementary Fig. S9). The 3-bp insertion, G58V, and V198M were all located outside the conserved zinc-finger and IGGH domains (Supplementary Fig. S8; Duan et al. 2019). Subcellular localization was examined for B843-type PgSHI1 (no coding variants, PgSHI1(B843)), and the cloned R102560-type *PgSHI1* allele carrying all three variants (PgSHI1(R102560)), each fused to GFP at the C-terminus. Both PgSHI1–GFP fusion proteins localized to the nucleus, consistent with the reported nuclear localization of OsSHI1 (Duan et al. 2019), and they showed no clear differences in GFP fluorescence intensity (Supplementary Fig. S10). MXfold2 prediction did not reveal an obvious alteration in local mRNA secondary structure around the variant sites between the B843-type and R102560-type *PgSHI1* sequences (Supplementary Fig. S11). AlphaFold 3 predicted similar overall structures for B843– and R102560-type PgSHI1, but an α-helix was predicted at the G58V site only in the R102560-type protein (Supplementary Fig. S12). Because OsSHI1 interacts with *Oryza sativa* IDEAL PLANT ARCHITECTURE1 (OsIPA1), a SQUAMOSA PROMOTER BINDING PROTEIN-LIKE (SPL)-family transcription factor involved in the regulation of plant architecture (Duan et al. 2019), we examined the predicted interaction of PgSHI1 with its putative pearl millet homolog, PgIPA1 (dpca4g302490.840). The PgIPA1–PgSHI1(B843) and PgIPA1–PgSHI1(R102560) complexes showed similar predicted interface areas and ΔiG values, with no substantial difference between the two PgSHI1 types (Supplementary Fig. S12 and Supplementary Table S6). Together, these results support the idea that *PgSHI1* is a candidate gene regulating culm height, although a clear functional effect of the associated CDS variants still remains unclear.

### *PgFY* is a candidate for a gene regulating days to panicle emergence

For days to panicle emergence, two associated polymorphisms were detected at Chr6:1,574,833 (*P* = 1.24e-6) and Chr6:1,940,237 (*P* = 4.15e-6) (Fig. 3a and 3b). An independent GWAS using phenotypic data from a 2023 field trial (Supplementary Table S2) identified additional associated polymorphisms in the same region at Chr6:1,574,224 (*P* = 1.83e-6) and Chr6:1,582,512 (*P* = 2.01e-6) (Supplementary Fig. S13). These four polymorphisms were located near or within *dpca6g389000.840*, with the associated polymorphism at Chr6:1,940,237 located in an intron (Fig. 3a, Fig. 3c and Supplementary Fig. S13a). *dpca6g389000.840* encodes a protein with high amino acid sequence similarity to Arabidopsis FY (AT5G13480), an RNA 3′-end-processing factor involved in flowering-time regulation (Simpson et al. 2003). Phylogenetic analysis placed dpca6g389000.840 with Arabidopsis FY and rice OsFY (LOC_Os01g72220; Lu et al. 2006) (Supplementary Fig. S14). GExA detected abundant *dpca6g389000.840* transcripts in vegetative and reproductive tissues, including stems and primary root tips (Supplementary Fig. S6). This observation is compatible with the reported expression of Arabidopsis *FY*, which is mainly expressed in shoot and root meristems and young tissues (Henderson et al. 2005). Based on these observations, *dpca6g389000.840* was designated *PgFY* and considered a candidate for a gene regulating days to panicle emergence.

Variants in the *PgFY* CDS were next investigated. None of the variants in the CDS represented in the GWAS marker set was significantly associated with days to panicle emergence. Reanalysis of the WGS data from the 21 lines identified three in-frame indel sites (Fig. 3c and Supplementary Fig. S15a). At Chr6:1,939,458, Hap. 2 carried a 6-bp insertion (GGCGGC), whereas Hap. 3 carried a 9-bp deletion (GGCGGCGGC) relative to Hap. 1 (Fig. 3c). At Chr6:1,939,401, Hap. 5 carried a 12-bp deletion (CAGCAGCAGCAA) relative to Hap. 4 (Supplementary Fig. S15a). At Chr6:1,939,571, Hap. 7, Hap. 8, and Hap. 9 carried 3-bp and 6-bp insertions and a 3-bp deletion, respectively, relative to Hap. 6 (Supplementary Fig. S15a).

To examine the relationship between these variants and days to panicle emergence within each GWAS lead-SNP genotype (Chr6:1,574,833) group, the 21 lines were classified according to the lead-SNP genotype and CDS diplotype. At Chr6:1,939,458, in the A/A genotype group at the GWAS lead SNP, Hap. 2/Hap. 2 lines showed later panicle emergence than Hap. 1/Hap. 1 lines (Fig. 3d). Comparisons involving Hap. 3 were not performed because the Hap. 1/Hap. 3 and Hap. 3/Hap. 3 classes were each represented by only one line (Fig. 3d). At Chr6:1,939,401, Hap. 5/Hap. 5 lines showed later panicle emergence than Hap. 4/Hap. 4 lines within the A/A lead-SNP group, whereas no significant difference was detected within the G/G group (Supplementary Fig. S15b). At Chr6:1,939,571, several genotype–diplotype classes contained only one line; among the A/A lead-SNP lines, however, Hap. 7/Hap. 7 was associated with later panicle emergence than Hap. 6/Hap. 6 (Supplementary Fig. S15c). Together, these results indicate that Hap. 2 at Chr6:1,939,458 was associated with later panicle emergence in the A/A genotype group at the GWAS lead SNP.

The potential effects of these variants on predicted RNA secondary structure, expression level, and protein structure were next examined. B00222 was homozygous for the three indel-containing haplotypes (Hap. 2, Hap. 5, and Hap. 7) examined above, whereas B94333 carried the corresponding reference haplotypes at these sites (Hap. 1, Hap. 4, and Hap. 6) (Supplementary Fig. S16). The predicted mRNA secondary structures showed no marked differences between the B00222-type and B94333-type *PgFY* mRNAs (Supplementary Fig. S17), and their predicted three-dimensional protein structures were also similar (Supplementary Fig. S18). *PgFY* expression levels in stems did not differ significantly between B94333 and B00222 (Supplementary Fig. S9b). Because Arabidopsis FY and rice OsFY interact with FLOWERING CONTROL LOCUS A (FCA) and OsFCA, respectively (Simpson et al. 2003; Lu et al. 2006), the predicted interactions of the two PgFY types with the putative pearl millet FCA homolog PgFCA (dpca7g511740.840) were compared. The predicted PgFCA–PgFY complexes showed similar interface areas and ΔiG values between B94333 and B00222 (Supplementary Fig. S18 and Supplementary Table S6). Together, these results support *PgFY* as a candidate for a gene regulating days to panicle emergence, although the examined variants in the CDS showed no clear effects on the molecular features analyzed.

### *PgEPF2* is a candidate for a gene regulating awn growth

For awn growth, a polymorphism associated with the trait was detected at Chr3:317,181,092 (*P* = 4.00e-9) (Fig. 4a and 4b). This polymorphism was located in the proximity of *dpca3g255860.840*, which encodes a protein similar to rice EPIDERMAL PATTERNING FACTOR 2 (OsEPF2; LOC_Os04g54490), an EPF/EPFL-family peptide involved in awn development in rice (Xiong et al. 2022). Another rice EPF/EPFL-family member, RAE2/OsEPFL1, also promotes awn elongation, and loss of RAE2 function contributes to awnlessness in cultivated Asian rice (Bessho-Uehara et al. 2016). Phylogenetic analysis showed that dpca3g255860.840 clustered with OsEPF2 within the EPF family (Supplementary Fig. S19). GExA detected abundant *dpca3g255860.840* transcripts in samples from multiple vegetative and reproductive tissues, including florets and tillering tissues (Supplementary Fig. S6). Its detection in reproductive tissues is compatible with the reported expression of *OsEPF2* in young panicles in rice (Xiong et al. 2022). Based on these observations, *dpca3g255860.840* was designated *PgEPF2* and considered a candidate for a gene regulating awn growth in pearl millet.

Variants in the *PgEPF2* CDS were next examined. None of the variants in the CDS represented in the GWAS marker set was significantly associated with awn growth. Reanalysis of the WGS data from the 21 lines identified a missense variant at Chr3:317,431,304 causing a Met57Thr substitution (M57T; Fig. 4c). M57T was located within the putative propeptide region, between the signal peptide cleavage site predicted by SignalP and the reported proteolytic cleavage sites of EPF2 and RAE2 (Supplementary Figs. S20 and S21; Engineer et al. 2014; Bessho-Uehara et al. 2016). When the 21 lines were classified according to the GWAS lead-SNP genotype (Chr3:317,181,092) and the *PgEPF2* CDS diplotype, awn growth did not differ significantly between Hap. 1/Hap. 1 and Hap. 2/Hap. 2 lines within either the G/G or A/A lead-SNP genotype group (Fig. 4d). Thus, *PgEPF2* is a candidate for a gene regulating awn growth based on its genomic position and homology to *OsEPF2*, whereas no significant association between the M57T variant and awn growth was detected in the 21 lines.

## Discussion

Using WGS-derived genotype data from 21 lines as a reference panel, genotype imputation substantially increased the completeness of the RAD-seq-derived genotype dataset and was accompanied by more localized GWAS signals in pearl millet (Table 1 and Fig. 1). Notably, significant associations were detected for traits that had not shown comparably strong signals in our previous GWAS using the RAD-seq-derived genotype dataset (Kambara et al. 2024). These included days to panicle emergence and awn growth, for which candidate genes were further investigated in the present study (Figs. 3 and 4). The more localized GWAS signals observed after imputation are consistent with previous findings that genotype imputation can increase marker density and facilitate association mapping in rice (Wang et al. 2018). Together, these observations support the utility of reference-panel-based genotype imputation for improving GWAS resolution in pearl millet.

In pearl millet, neither reference-panel-based genotype imputation nor the establishment of a reference panel for GWAS has previously been reported. In the present study, a reference panel comprising 21 B– and R-lines was used to impute genotypes in 81 lines. B– and R-lines play central roles in pearl millet hybrid breeding as maintainer and restorer lines, respectively, and considerable genetic diversity has been reported within both groups (Mula et al. 2007; Ramya et al. 2018). In this study, the leave-one-out validation yielded a mean genotype concordance of 89.9% (Supplementary Table S3 and Supplementary Fig. S2), suggesting that the 21-line reference panel was useful for genotype imputation in the B– and R-line germplasm examined here. Reference panels of various sizes have been used for genotype imputation in other crops. In rice, a reference panel comprising 3,023 accessions was used to impute genotypes in 1,568 samples (Wang et al. 2018). In contrast, a reference panel comprising 62 lines was used to impute genotypes in 307 accessions in wheat, improving the resolution of GWAS (Nyine et al. 2019). In soybean, a reference panel comprising 25 parental lines was used to impute genotypes in 2,669 recombinant inbred lines, improving the detection of association signals in GWAS (Kim et al. 2026). These studies suggest that reference panels comprising a few dozen lines can be useful for genotype imputation in larger target populations. The 21-line reference panel established in the present study may also be useful for genotype imputation in larger collections of pearl millet lines beyond the 81 lines imputed here, although its applicability to populations with different genetic backgrounds remains to be evaluated.

Among the associations detected by GWAS using the imputed genotype dataset, the chromosome 4 association for culm height implicated *PgSHI1* as a candidate gene (Fig. 2a). Overexpression of *OsSHI1* reduces plant height in rice (Duan et al. 2019), supporting the potential involvement of *PgSHI1* in regulating culm height in pearl millet. Reanalysis of the WGS data identified a 3-bp in-frame insertion in the *PgSHI1* CDS, which was associated with differences in culm height among the 21 lines. *PgSHI1* transcript abundance in stem tissue did not differ significantly between B843 and R102560, and the two PgSHI1–GFP fusion proteins showed similar nuclear localization. Predicted mRNA secondary structures and overall protein structures were also similar, although an α-helix was predicted around the G58V site only in the R102560-type PgSHI1 protein (Supplementary Fig. S12). In addition, no substantial difference was predicted in the interaction of the PgSHI1 proteins with PgIPA1. These analyses did not reveal clear effects of the variants on transcript abundance, subcellular localization, predicted structure, or the examined protein interaction. Functional validation of individual *PgSHI1* variants will therefore be required to determine whether they contribute to the culm height variation. The chromosome 6 association for days to panicle emergence implicated *PgFY* as a candidate gene (Fig. 3a). An independent GWAS using phenotypic data obtained in 2023 detected an associated polymorphism at nearly the same position as the strongest signal identified using the phenotypic dataset from Kambara et al. (2024) (Supplementary Fig. S13). The recurrence of the chromosome 6 association across the two phenotypic datasets provides independent support for a locus affecting days to panicle emergence. FY is involved in RNA 3′-end processing and flowering-time regulation in Arabidopsis, and hypomorphic *fy* alleles exhibit late flowering (Simpson et al. 2003; Feng et al. 2011). These observations support *PgFY* as a candidate for regulating panicle emergence in pearl millet. Several in-frame indels were identified within the *PgFY* CDS (Fig. 3c and Supplementary Fig. S15a). At Chr6:1,939,458, lines homozygous for the 6-bp insertion showed later panicle emergence than lines without the insertion within the A/A genotype group at the GWAS lead SNP. These results raise the possibility that the 6-bp insertion contributes to delayed panicle emergence in pearl millet. However, no marked differences were detected between the examined *PgFY* alleles in predicted mRNA secondary structure, overall protein structure, or predicted interaction with PgFCA (Supplementary Figs. S17 and S18). Thus, as for *PgSHI1*, the present analyses support an association between a coding haplotype and the phenotype but did not reveal its molecular effect.

Among the CDS variants associated with phenotypic differences in *PgSHI1* and *PgFY* were in-frame insertions encoding glycine residues near the N terminus (Figs. 2c and 3c). These insertions introduced GC-rich glycine codons in the N-terminal coding regions, altering both the local nucleotide composition and the encoded amino-acid sequences. In *Escherichia coli*, increased GC content and stable mRNA secondary structure near the start codon can reduce protein production by limiting translation initiation (Kudla et al. 2009; Bentele et al. 2013). Although MXfold2 did not predict marked differences in mRNA secondary structure among the alleles (Supplementary Figs. S11 and S17), such predictions may not fully capture effects on translation in vivo. More broadly, codon usage can influence gene expression (Tsugama and Kambara 2026), raising the possibility that the GC-rich glycine codons introduced by these insertions affect protein production through codon-dependent mechanisms. Changes involving N-terminal Gly residues have also been reported to alter protein production. In mammalian cells, insertion of a single Gly immediately after the initiating Met markedly reduced the expression of a luciferase reporter containing the N-terminal sequence of extracellular signal-regulated kinase 2 (ERK2) (Buscà et al. 2023). Similarly, replacement of Gly at the fourth residue of superfolder green fluorescent protein (sfGFP) altered protein synthesis in an *E. coli*-derived cell-free translation system (Fuse-Murakami et al. 2024). Although comparable effects have not been directly demonstrated in plants, the N-terminal insertions in PgSHI1 and PgFY may affect protein production through altered local RNA structure or translation efficiency. For PgSHI1, no obvious difference in GFP fluorescence was observed between the two PgSHI1–GFP fusion proteins (Supplementary Fig. S10). However, because this transient assay examined a GFP-fused protein rather than endogenous PgSHI1, it does not exclude an effect of the variants on PgSHI1 protein production in its native context. Direct quantification of endogenous PgSHI1 and PgFY protein abundance will be required to test this possibility.

The chromosome 3 association for awn growth implicated *PgEPF2* as a candidate gene (Fig. 4a). *PgEPF2* is homologous to rice *OsEPF2*, and loss of *OsEPF2* function reduces awn length in rice (Xiong et al. 2022), providing functional support for a role of *PgEPF2* in awn growth. An M57T substitution was identified in the putative propeptide region of PgEPF2 (Fig. 4c and Supplementary Fig. S20). Awn growth did not differ significantly between the *PgEPF2* CDS diplotypes within either genotype group at the GWAS lead SNP. Thus, the M57T variant was not significantly associated with awn growth in the 21 lines. The GWAS lead SNP is located approximately 250 kb from the *PgEPF2* CDS (Fig. 4a). The association may instead reflect a noncoding variant within the associated region that affects the spatial or temporal expression of *PgEPF2* rather than its protein sequence. Regulatory variants underlying agronomic traits have been reported in rice, including variants affecting *qSH1* and *FRIZZY PANICLE* (*FZP*) expression (Konishi et al. 2006; Huang et al. 2018). Analysis of *PgEPF2* expression in developing panicles and awn primordia, together with functional analysis of candidate noncoding variants, will be required to determine whether the association on chromosome 3 involves regulation of *PgEPF2*.

Overall, these results demonstrate that reference-panel-based genotype imputation can improve genotype completeness and GWAS resolution in pearl millet. The present analyses further support *PgSHI1*, *PgFY*, and *PgEPF2* as candidate genes for culm height, days to panicle emergence, and awn growth, respectively. These findings provide a foundation for further genetic dissection of agronomic traits and more effective utilization of existing genotype resources in future pearl millet research and breeding.

## Author Contributions

All authors contributed to the study conception and design. Material preparation was performed by S.K.G. Data collection and analysis were performed by K.K. and D.T. The first draft of the manuscript was written by K.K., and all authors commented on previous versions of the manuscript. All authors read and approved the final manuscript.

## Acknowledgments

We are grateful to ICRISAT for providing the pearl millet seeds for this study. We thank the technical staff of the Institute for Sustainable Agro-ecosystem Services, The University of Tokyo, for preparing the fields used in this study.

## Statements and Declarations

### Data availability

All data and analysis scripts are available in the figshare repository (Kambara and Tsugama 2026, https://doi.org/10.6084/m9.figshare.34419156). The WGS data generated in this study are available in the DNA Data Bank of Japan (DDBJ) Sequence Read Archive (DRA) under BioProject accession number PRJDB45871.

