## Supplementary Figures for "Genotype imputation improves GWAS resolution and facilitates candidate gene identification in pearl millet"

**Title**

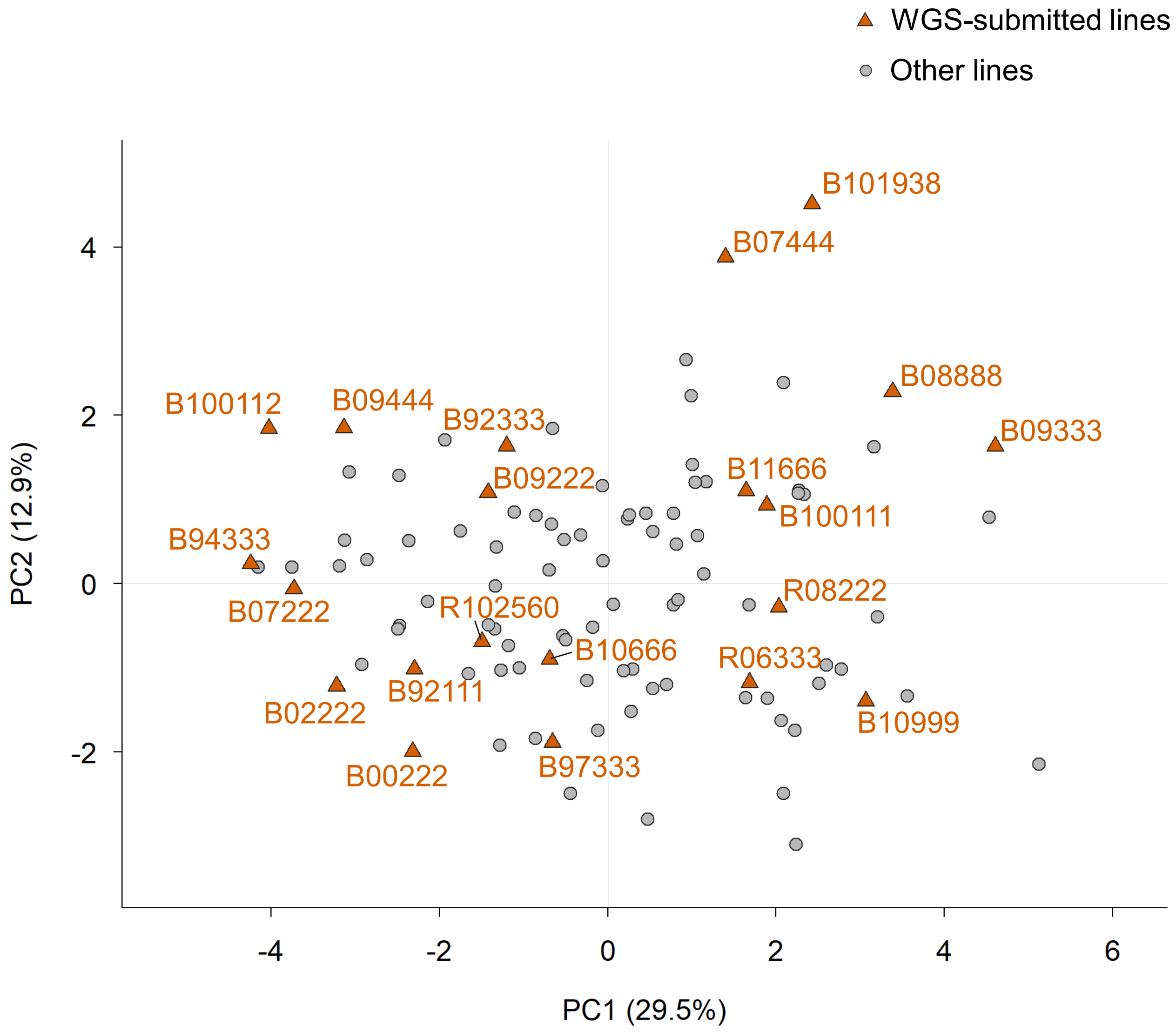


**Supplementary Fig. S1 Principal component analysis of phenotypic variation among pearl millet elite lines**

Principal component analysis (PCA) was performed for 101 lines using 15 of the 16 phenotypic traits reported by Kambara et al. (2024), excluding “pigmentation of nodes”. Each point represents an individual line. PC1 and PC2 explained 29.5% and 12.9% of the total phenotypic variance, respectively.

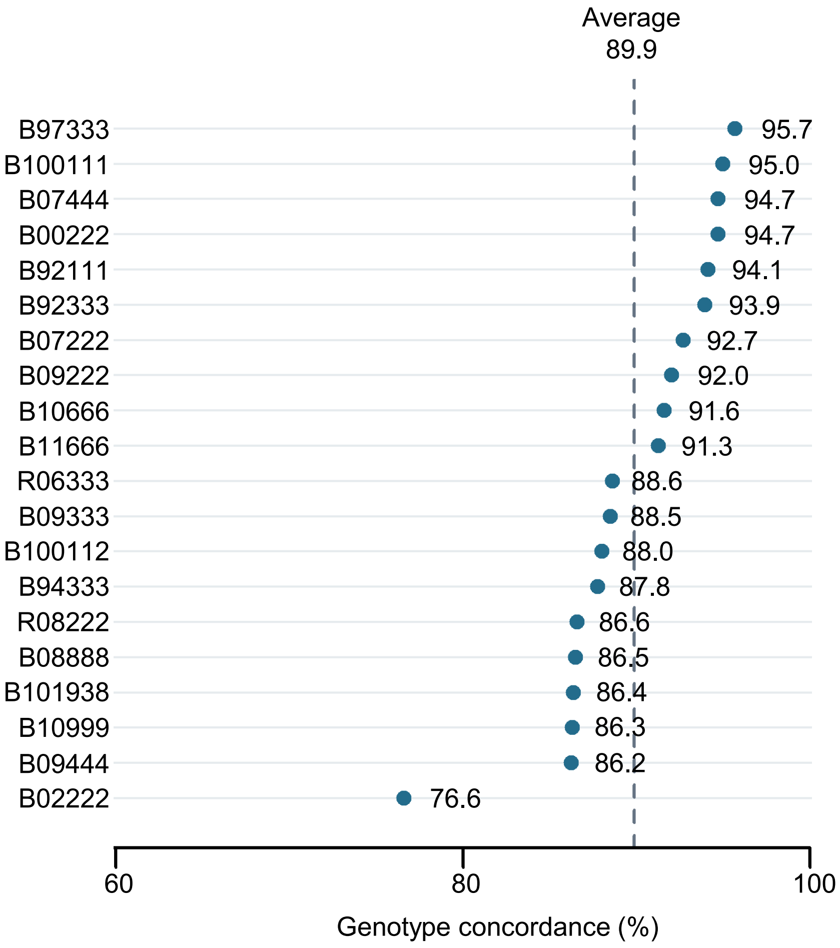


**Supplementary Fig. S2 Leave-one-out validation of genotype imputation accuracy**

Genotype concordance between imputed and WGS-derived genotypes was evaluated for 20 reference-panel lines by leave-one-out validation. For each line, the line was excluded from the reference panel and its RAD-seq-derived genotypes were imputed using the remaining 20 reference lines. Each point represents the genotype concordance for one line, and the values shown indicate concordance percentages. The dashed vertical line indicates the mean concordance across the 20 lines (89.9%).

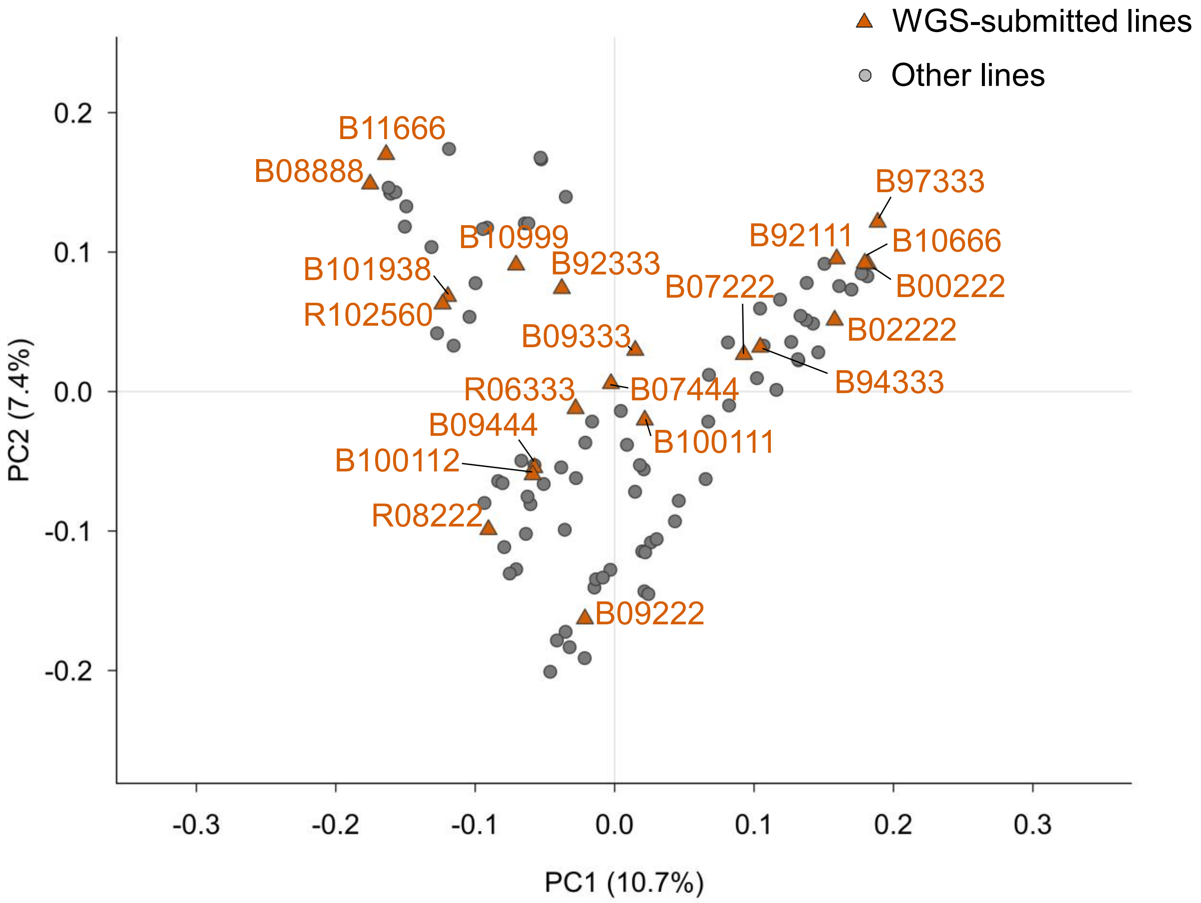


**Supplementary Fig. S3 Principal component analysis of genome-wide genetic variation among pearl millet elite lines**

Principal component analysis (PCA) was performed for 102 lines using 2,153,178 biallelic SNPs. Each point represents an individual line. PC1 and PC2 explained 10.7% and 7.4% of the total genetic variation, respectively.

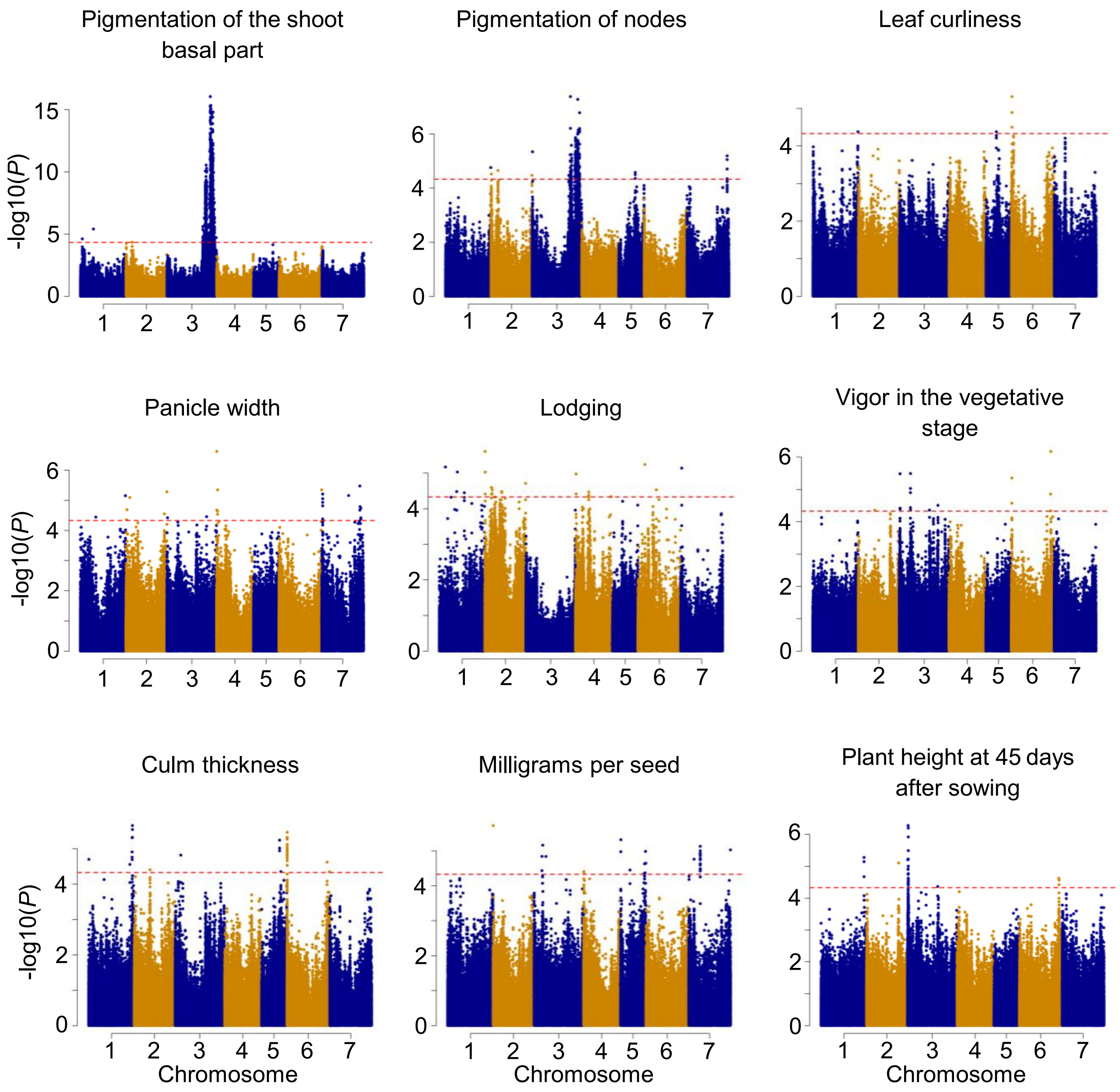


**Supplementary Fig. S4 Manhattan plots for additional traits using the imputed genotype dataset**

Manhattan plots for pigmentation of the shoot basal part, pigmentation of nodes, leaf curliness, panicle width, lodging, vigor at the vegetative stage, culm thickness, milligrams per seed, and plant height at 45 days after sowing. The x-axis represents chromosome position, and the y-axis represents $-\log_{10}(P)$. Red dashed lines indicate the genome-wide significance threshold ($P$ < 4.66e-5), determined based on the effective number of independent tests using the method of Li and Ji (2005).

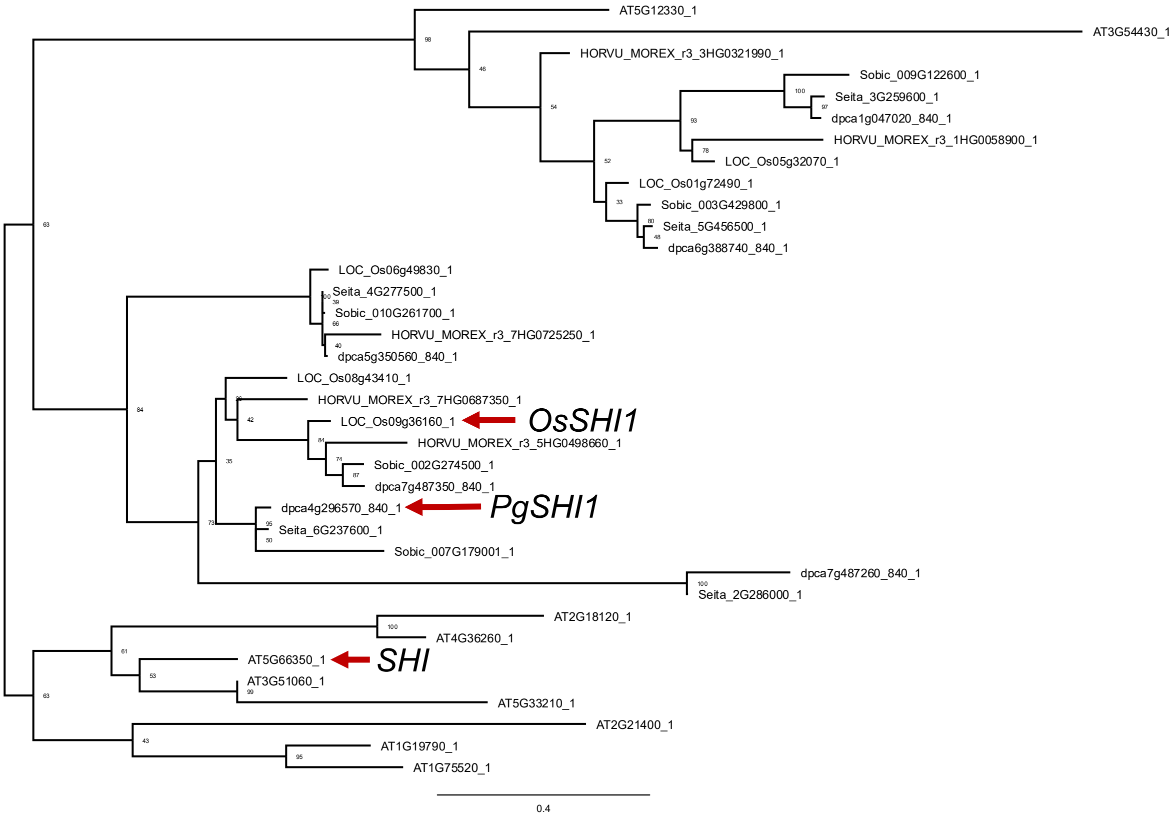


**Supplementary Fig. S5 Maximum-likelihood phylogenetic tree of *PgSHI1* homologs**

Numbers at nodes indicate bootstrap support values (%) based on 1,000 replicates. Red arrows indicate *OsSHI1* (*LOC_Os09g36160*), *PgSHI1* (*dpca4g296570.840*) and *SHI* (*AT5G66350*).

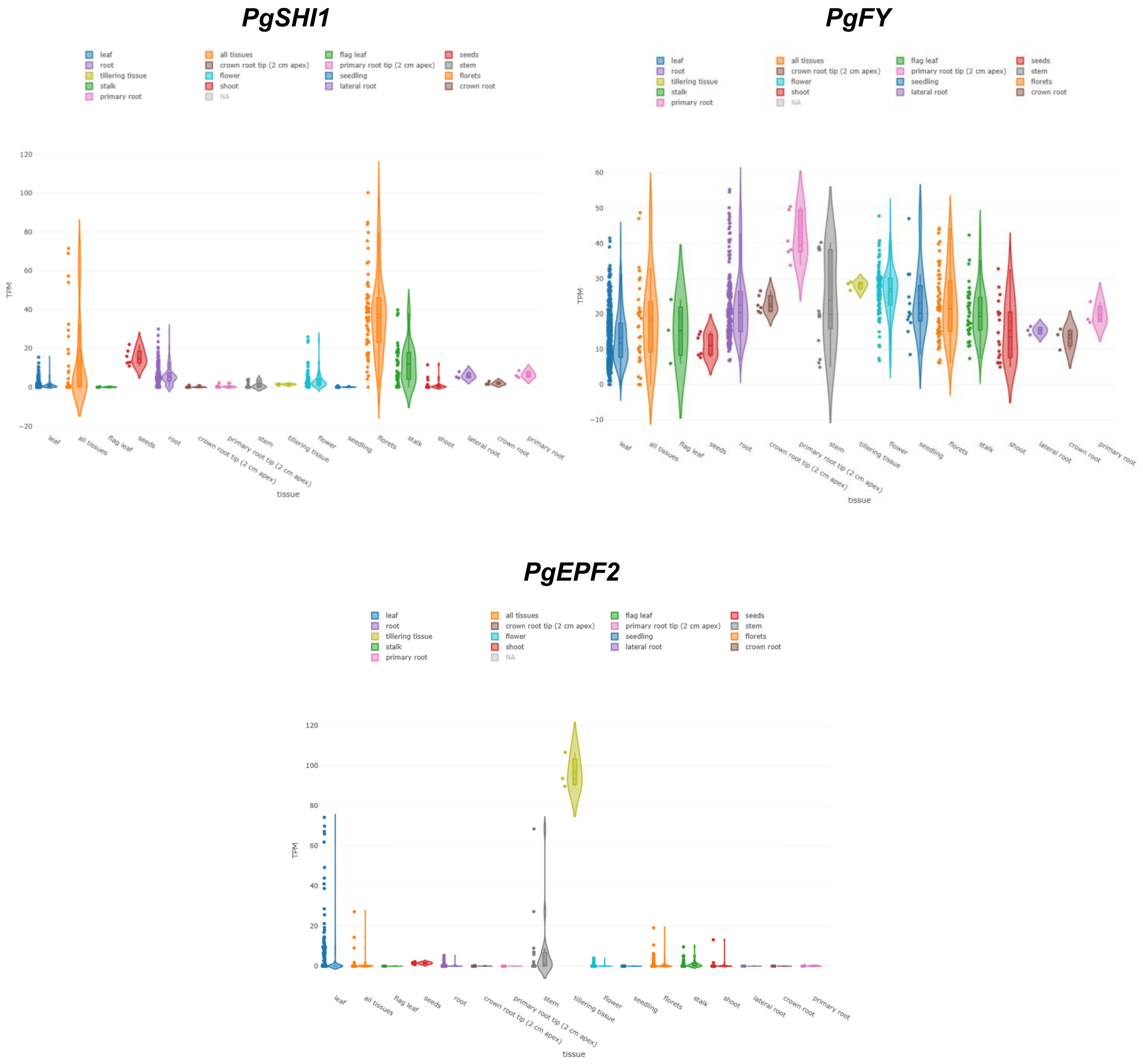


**Supplementary Fig. S6 Expression profiles of candidate genes in pearl millet tissues**
Expression profiles of *PgSHI1* (*dpca4g296570.840*), *PgFY* (*dpca6g389000.840*), and *PgEPF2* (*dpca3g255860.840*) in the Grass Expression Atlas (GExA). Transcript abundance is shown as transcripts per million (TPM) across tissue categories. Individual points represent RNA-seq samples.

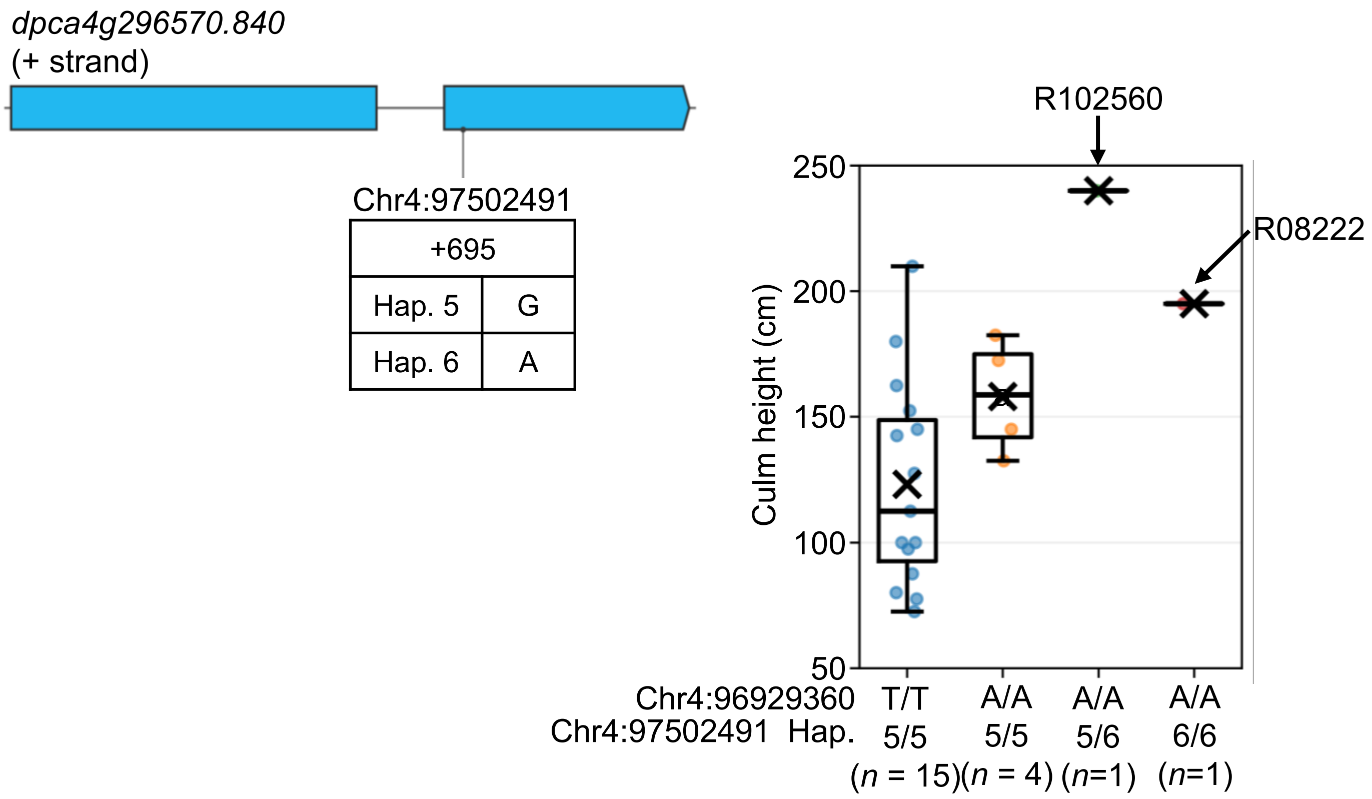


**Supplementary Fig. S7 A missense variant site at Chr4:97,502,491 causing a Val198Met substitution in PgSHI1**
Schematic representation of *PgSHI1* and sequence variants at Chr4:97,502,491. Boxes indicate exons, with untranslated regions (UTRs) shown in white and coding regions shown in blue; black lines indicate introns. Culm height of the 21 lines classified by genotype at the GWAS lead SNP Chr4:96,929,360 and diplotype at Chr4:97,502,491. Diplotypes at Chr4:97,502,491 were determined by reanalysis of the WGS data from the 21 lines. Individual points represent individual lines; crosses indicate means, horizontal lines indicate medians, and boxes indicate interquartile ranges. Sample sizes (*n*) are indicated below each group.

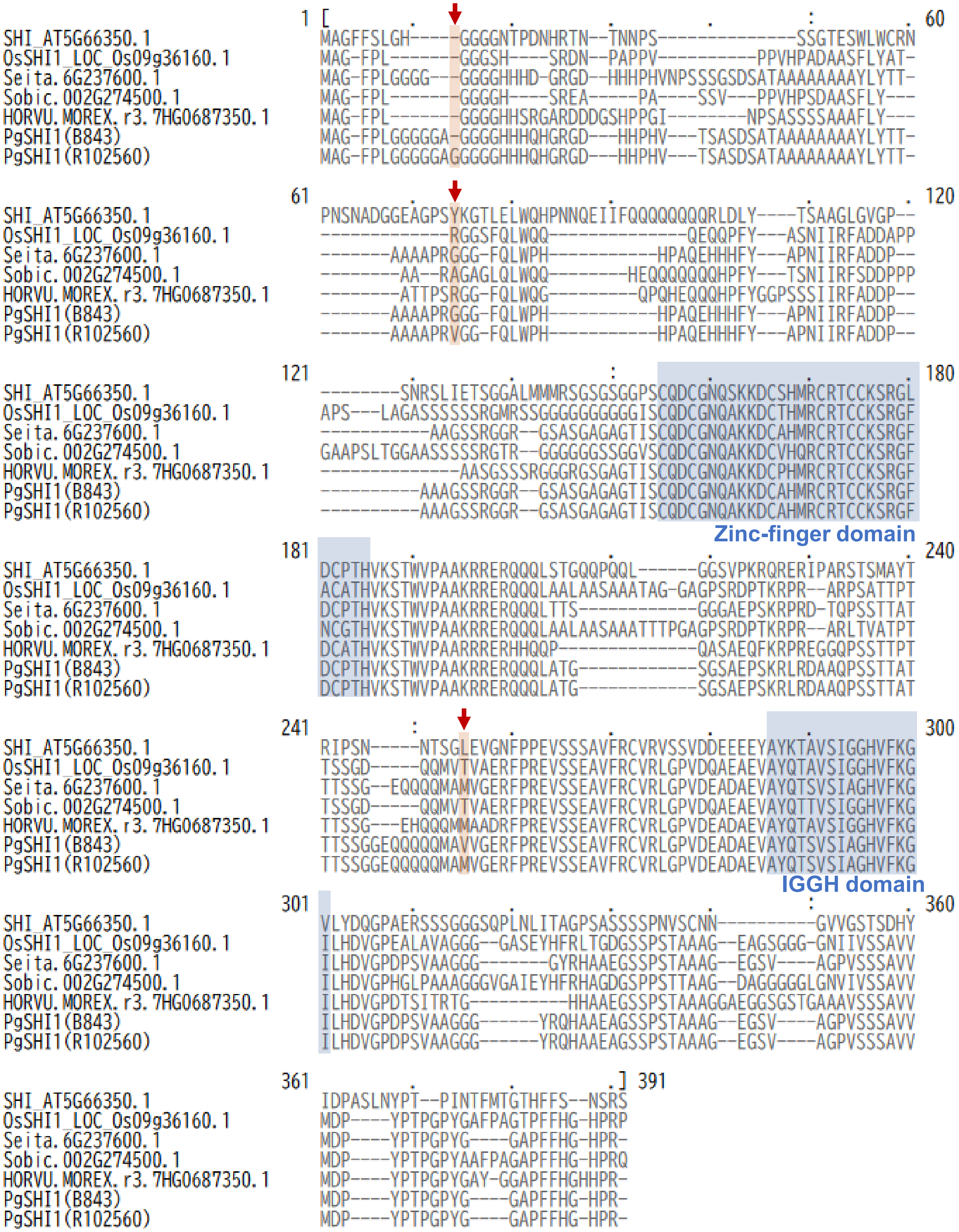


**Supplementary Fig. S8 Multiple sequence alignment of SHI family proteins and the positions of coding variants in PgSHI1**

Amino acid sequences of Arabidopsis SHI (AT5G66350.1), rice OsSHI1 (LOC_Os09g36160.1), and SHI homologs from *Setaria italica*, *Sorghum bicolor*, and *Hordeum vulgare*, together with PgSHI1 sequences from B843 and R102560, were aligned using MEGA11 (Tamura et al. 2021). B843 represents the PgSHI1 allele without the three coding variants examined in this study, whereas R102560 carries the 3-bp insertion and the G58V and V198M substitutions. Red arrows indicate the positions of these three variants. The zinc-finger and IGGH domains are shaded in blue, with their positions defined according to Duan et al. (2019).

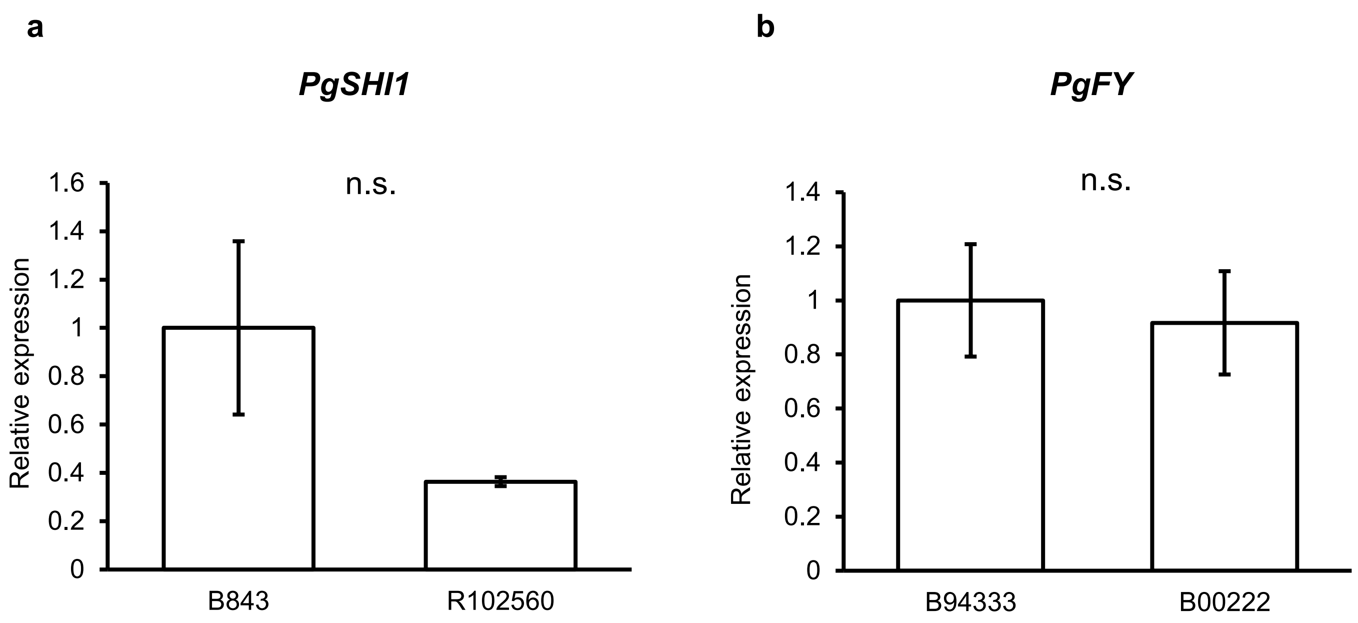


**Supplementary Fig. S9 Relative expression levels of *PgSHI1* and *PgFY* determined by qRT-PCR**

Relative expression levels of **a** *PgSHI1* in B843 and R102560 and **b** *PgFY* in B94333 and B00222. Expression levels were calculated using the comparative Ct (ΔΔCt) method with *PgEF1α* as the internal control. Data were obtained from three biological replicates ($n = 3$). Statistical significance was assessed using Welch's *t*-test ($P < 0.05$). n.s., not significant. Bars represent means, and error bars indicate SD.

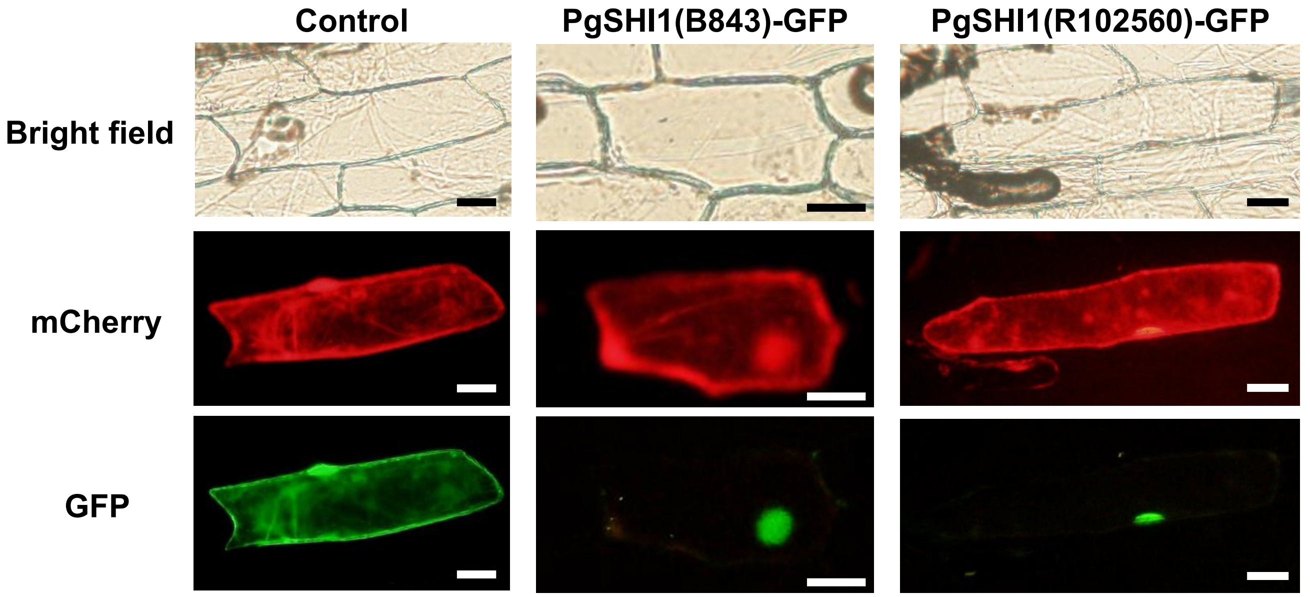


**Supplementary Fig. S10 Subcellular localization of B843- and R102560-type PgSHI1**
PgSHI1(B843)-GFP and PgSHI1(R102560)-GFP were transiently expressed in onion epidermal cells together with mCherry. GFP alone was used as the control. Bright-field, mCherry, and GFP fluorescence images are shown. Scale bars = 50 µm.

B843 R102560

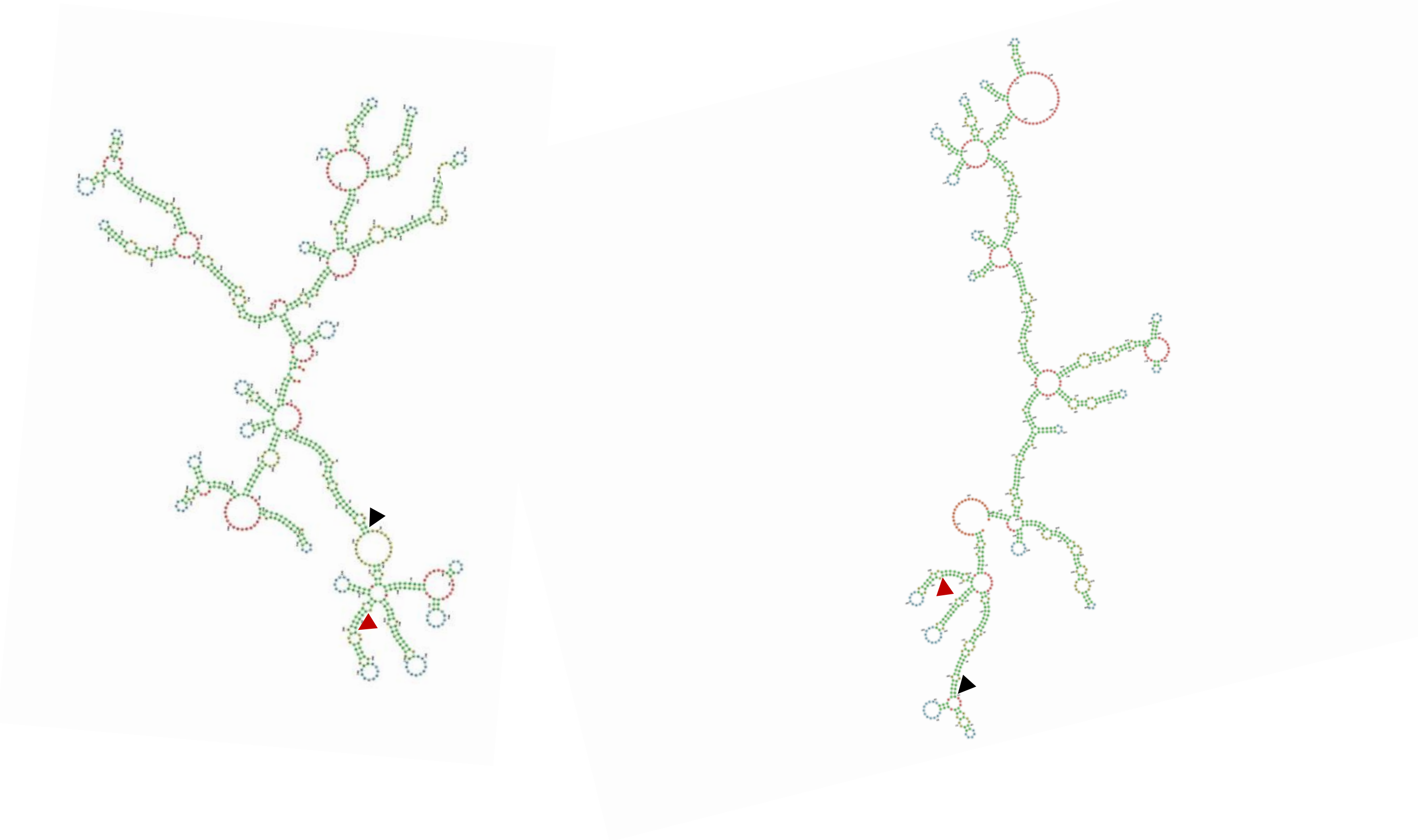

**Supplementary Fig. S11 Predicted RNA secondary structures of B843- and R102560-type *PgSHI1***

RNA secondary structures of the *PgSHI1* coding sequences from B843 and R102560 were predicted using MXfold2. B843 carries neither the 3-bp GGA insertion nor the G58V substitution, whereas R102560 carries both variants. Red and black arrowheads indicate the positions of the 3-bp GGA insertion and the G-to-T substitution causing G58V, respectively.

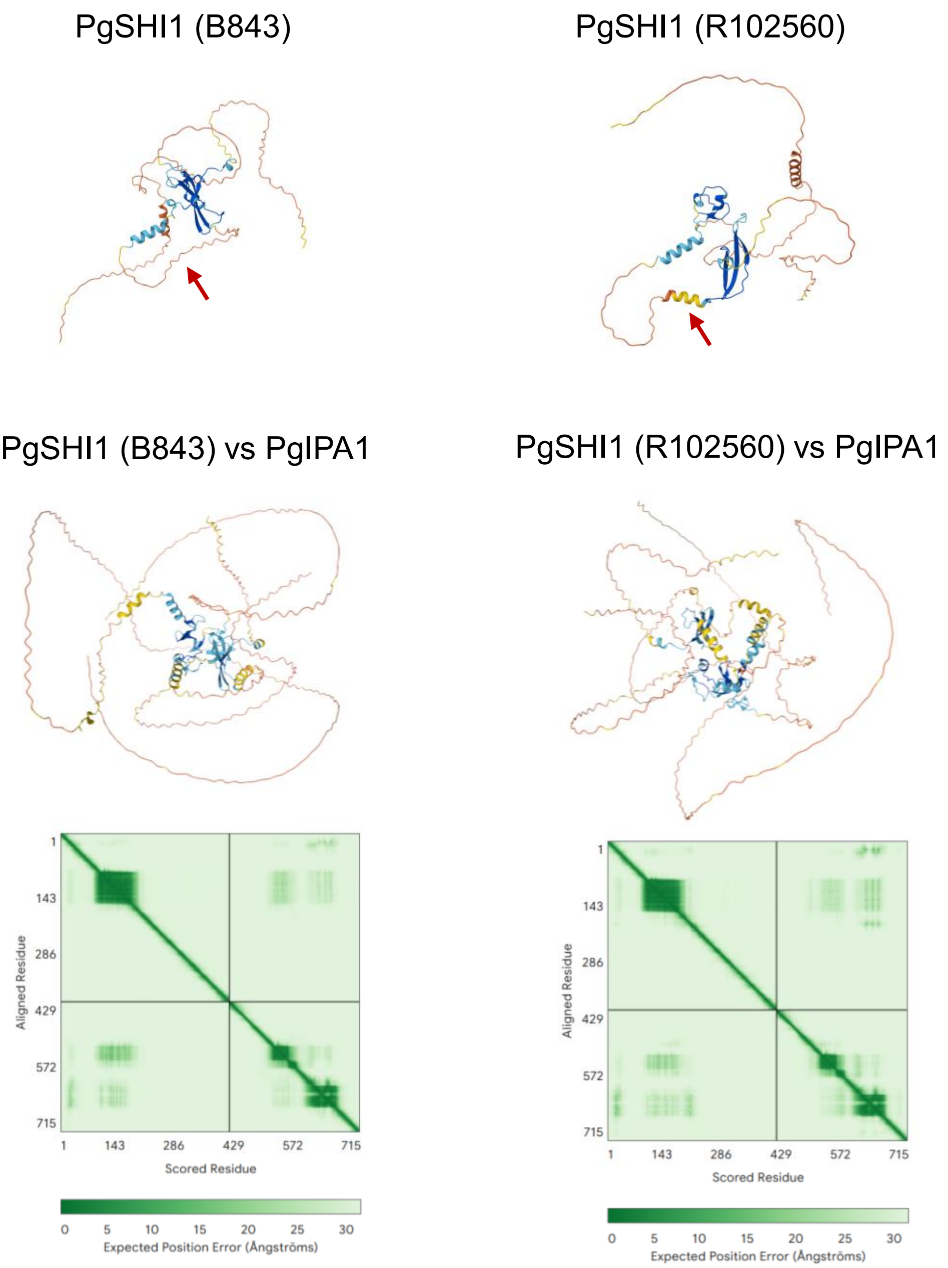


**Supplementary Fig. S12 Predicted structures of B843- and R102560-type PgSHI1 and their complexes with PgIPA1**

Three-dimensional structures of B843- and R102560-type PgSHI1 and their complexes with PgIPA1 were predicted using AlphaFold 3. Red arrows indicate the position of the G58V substitution. PgIPA1 (dpca4g302490.840) was identified as a putative pearl millet homolog of rice OsIPA1 (LOC_Os08g39890) by BLASTP analysis. Predicted aligned error (PAE) plots for the PgSHI1–PgIPA1 complexes are shown below the corresponding structural models.

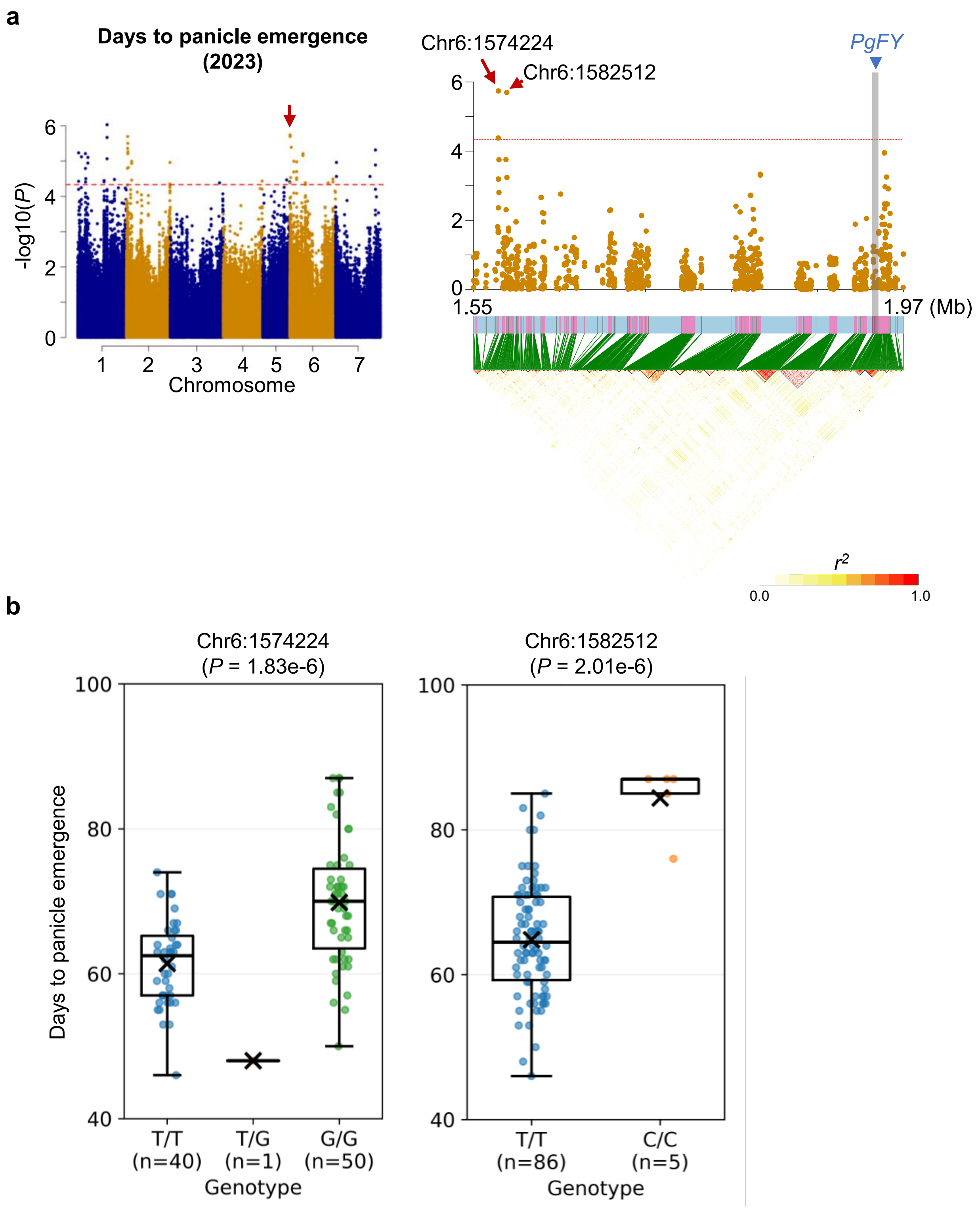


**Supplementary Fig. S13 GWAS for days to panicle emergence using phenotypic data from the 2023 field trial**

**a** Manhattan plot for days to panicle emergence measured in the 2023 field trial (left) and regional association plot with linkage disequilibrium (LD) around the chromosome 6 association peak (right). The red dashed line indicates the genome-wide significance threshold ($P < 4.66e\text{-}5$). The red arrows indicate the associated SNPs at Chr6:1,574,224 and Chr6:1,582,512, and the blue triangle indicates the position of *PgFY* (*dpca6g389000.840*). Pairwise LD is shown as $r^2$. **b** Days to panicle emergence of lines classified by genotype at Chr6:1,574,224 and Chr6:1,582,512. The phenotype and genotype data are those used in the GWAS shown in **a**. Individual points represent individual lines; crosses indicate means, horizontal lines indicate medians, and boxes indicate interquartile ranges. Sample sizes (*n*) are indicated below each group.

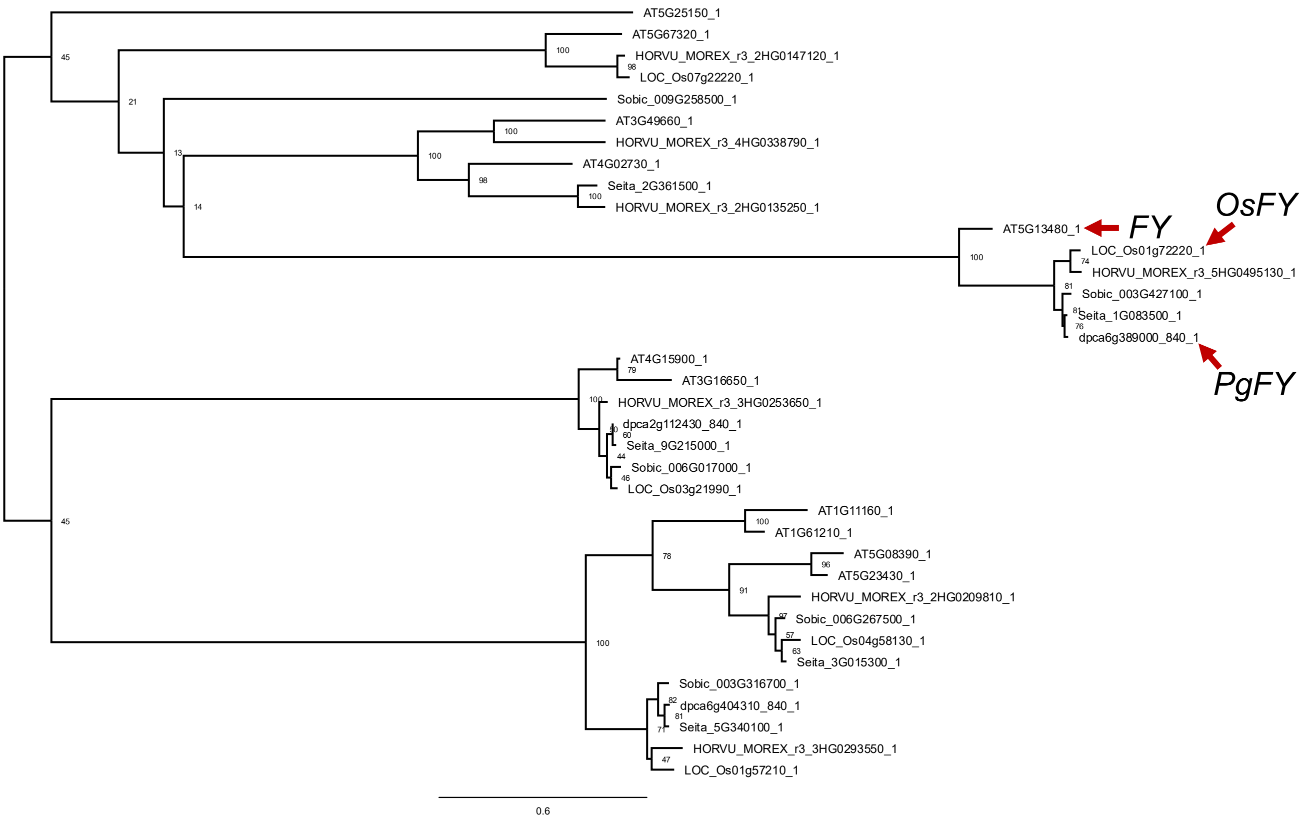


**Supplementary Fig. S14 Maximum-likelihood phylogenetic tree of *PgFY* homologs**

Numbers at nodes indicate bootstrap support values (%) based on 1,000 replicates. Red arrows indicate *OsFY* (*LOC_Os01g72220*), *PgFY* (*dpca6g389000.840*) and *FY* (*AT5G13480*).

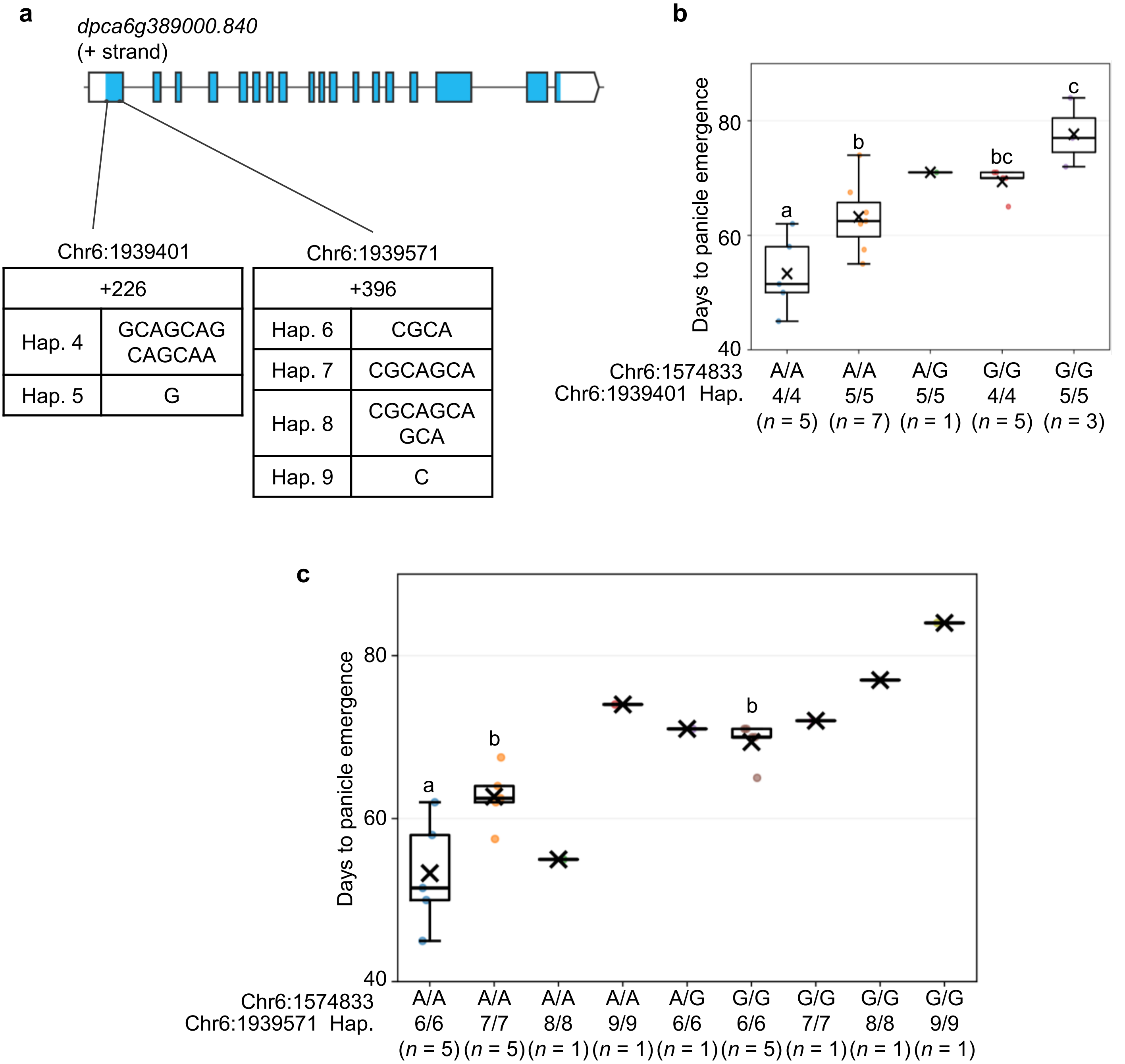


**Supplementary Fig. S15 Relationship between additional *PgFY* coding indels and days to panicle emergence**

**a** Schematic representation of *PgFY* and sequence variants at Chr6:1,939,401 and Chr6:1,939,571. Boxes indicate exons, with untranslated regions (UTRs) shown in white and coding regions shown in blue; black lines indicate introns. b Days to panicle emergence of the 21 lines classified by genotype at Chr6:1,574,833 and diplotype at Chr6:1,939,401. Hap. 5 carries a 12-bp deletion relative to Hap. 4. **c** Days to panicle emergence of the 21 lines classified by genotype at Chr6:1,574,833 and diplotype at Chr6:1,939,571. At Chr6:1,939,571, Hap. 7 and Hap. 8 carry 3-bp and 6-bp insertions, respectively, whereas Hap. 9 carries a 3-bp deletion relative to Hap. 6. Diplotypes at Chr6:1,939,401 and Chr6:1,939,571 were determined by reanalysis of the WGS data from the 21 lines. Individual points represent individual lines; crosses indicate means, horizontal lines indicate medians, and boxes indicate interquartile ranges. Sample sizes (*n*) are indicated below each group. Different lowercase letters indicate significant differences among groups ($P < 0.05$; Tukey–Kramer test). Groups represented by a single line were excluded from statistical comparisons.

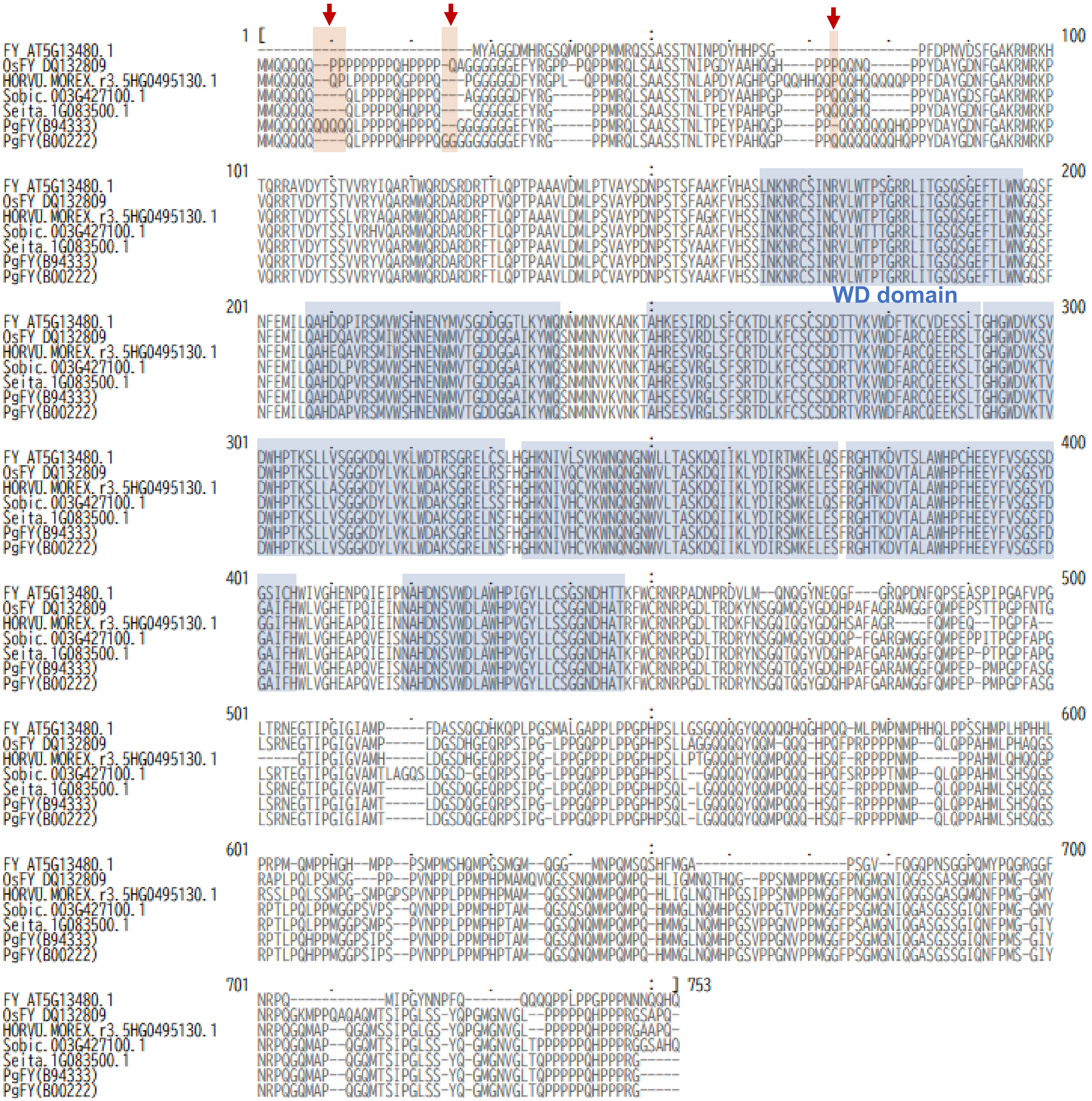

**Supplementary Fig. S16 Multiple sequence alignment of FY proteins and the positions of coding indels in PgFY**

Amino acid sequences of Arabidopsis FY (AT5G13480.1), rice OsFY, and FY homologs from *Hordeum vulgare*, *Sorghum bicolor*, and *Setaria italica*, together with PgFY sequences from B94333 and B00222, were aligned using MEGA11. B94333 carries the reference haplotypes at the three *PgFY* indel sites examined, whereas B00222 is homozygous for the corresponding indel-containing haplotypes. Red arrows indicate the positions of the three indel sites. The WD domain is shaded in blue, with its position defined according to Lu et al. (2006). The OsFY amino acid sequence was translated from the cDNA sequence reported by Lu et al. (2006) (GenBank accession DQ132809).

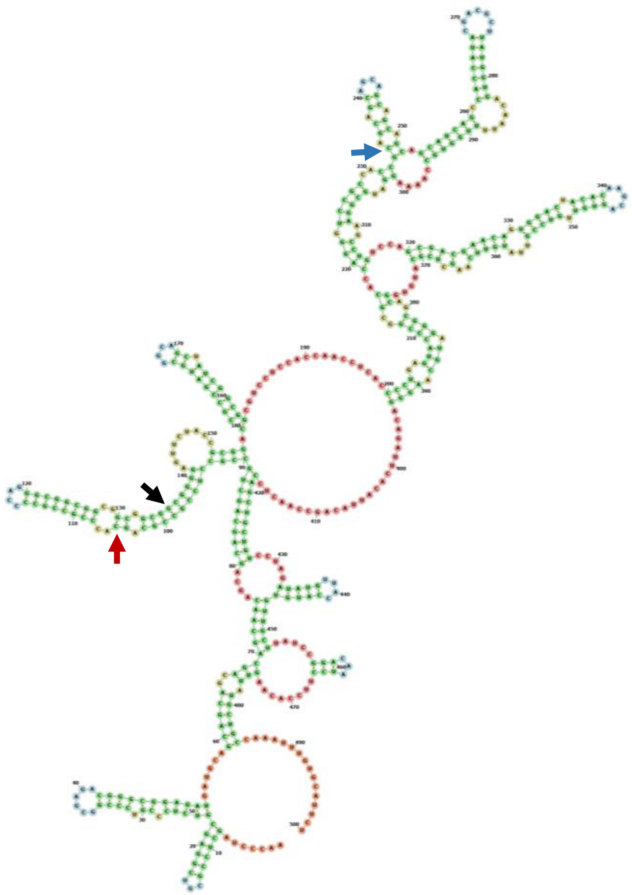

*PgFY* (B94333)

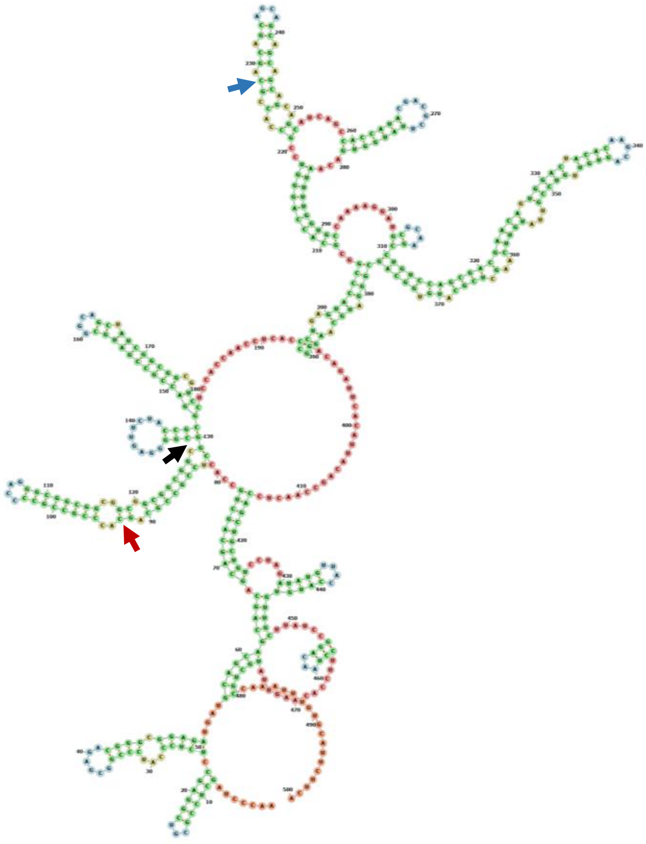

*PgFY* (B00222)

**Supplementary Fig. S17 Predicted RNA secondary structures of B94333- and B00222-type *PgFY***

RNA secondary structures of the predicted 500-nt *PgFY* transcript regions spanning from 50 nt upstream to 450 nt downstream of the start codon were predicted using MXfold2. B94333 carries the reference haplotypes at the three indel sites examined, whereas B00222 is homozygous for the corresponding indel-containing haplotypes. Colored arrowheads indicate the positions of the three indel sites.

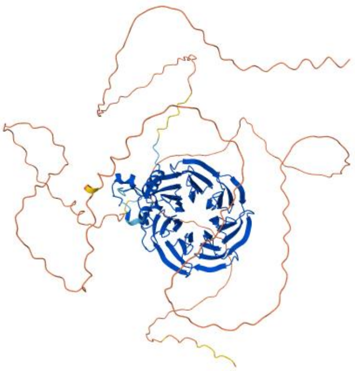

PgFY (B94333)

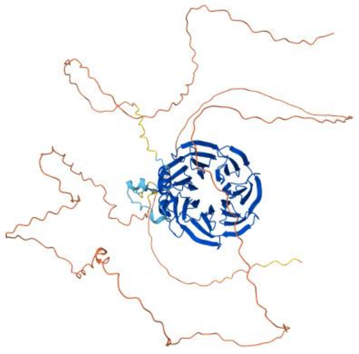

PgFY (B00222)

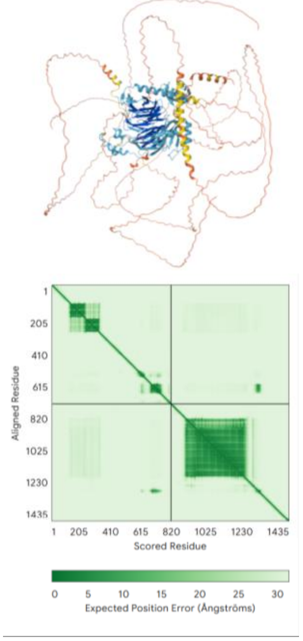


PgFY (B94333) vs PgFCA

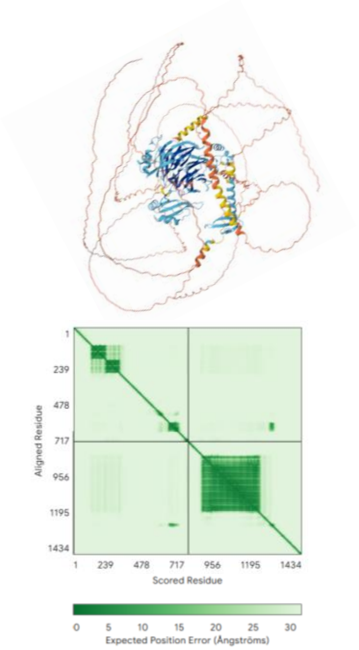


PgFY (B00222) vs PgFCA

**Supplementary Fig. S18 Predicted structures of B94333- and B00222-type PgFY and their complexes with PgFCA**

Three-dimensional structures of B94333- and B00222-type PgFY and their complexes with PgFCA were predicted using AlphaFold 3. B94333 carries the reference haplotypes at the three *PgFY* indel sites examined, whereas B00222 is homozygous for the corresponding indel-containing haplotypes. PgFCA (*dpca7g511740.840*) was identified as a putative pearl millet homolog of rice OsFCA (LOC_Os09g03610) by BLASTP analysis. Predicted aligned error (PAE) plots for the PgFY–PgFCA complexes are shown below the corresponding structural models.

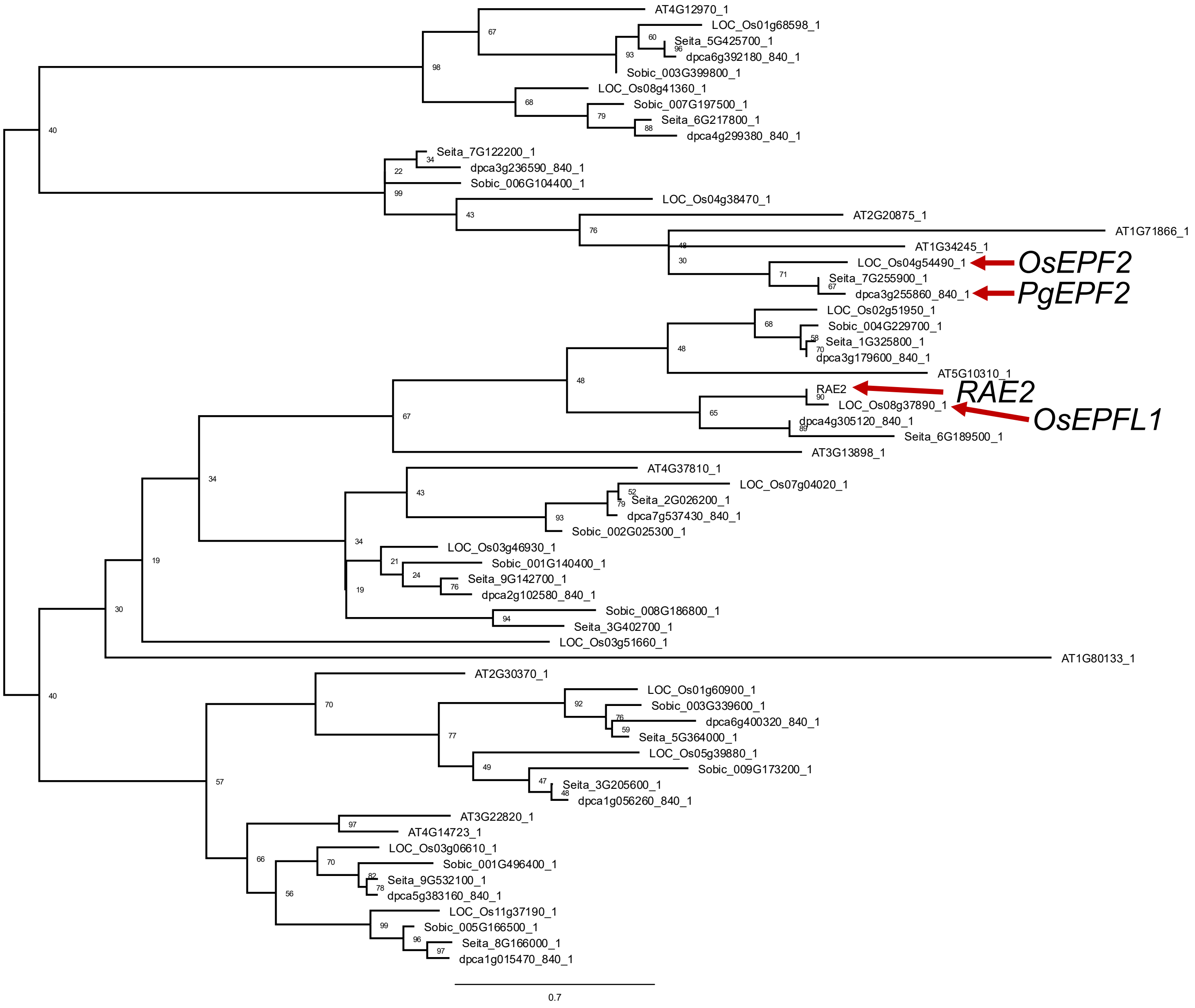


**Supplementary Fig. S19 Maximum-likelihood phylogenetic tree of EPF family proteins.** Numbers at nodes indicate bootstrap support values (%) from 1,000 replicates. Red arrows indicate *OsEPF2* (*LOC_Os04g54490*), *PgEPF2* (*dpca3g255860.840*)*, RAE2, and OsEPFL1* (*LOC_Os08g37890.1*)

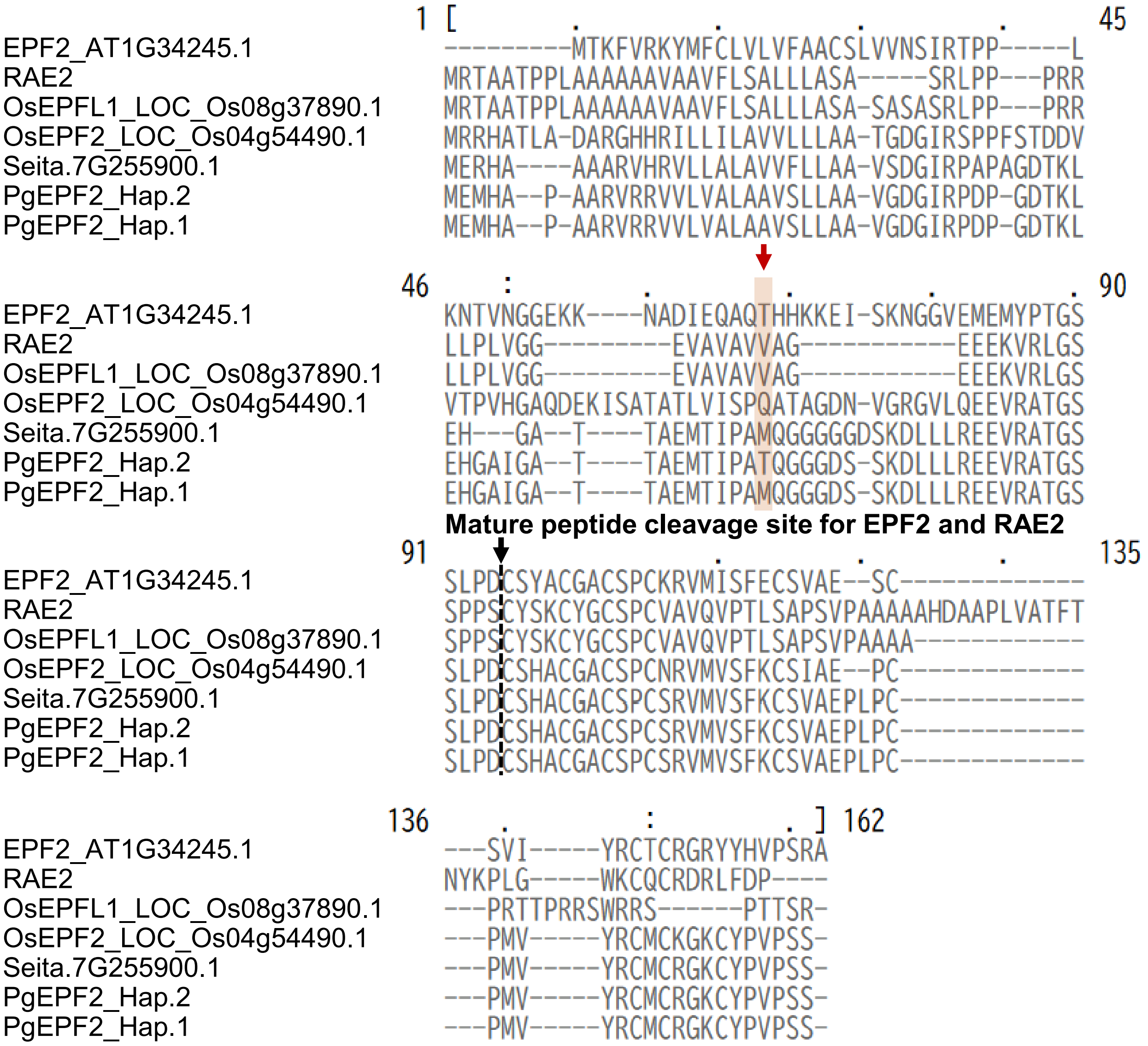


**Supplementary Fig. S20 Multiple sequence alignment of EPF/EPFL family proteins and the position of the M57T variant in PgEPF2**

Amino acid sequences of Arabidopsis EPF2 (AT1G34245.1), *Oryza glaberrima* RAE2, *O. sativa* OsEPFL1 (LOC_Os08g37890.1) and OsEPF2 (LOC_Os04g54490.1), *Setaria italica* 7G255900.1, and PgEPF2 Hap. 1 and Hap. 2 were aligned using MEGA11. The black arrow indicates the proteolytic cleavage sites associated with maturation of EPF2 and RAE2 reported by Engineer et al. (2014) and Bessho-Uehara et al. (2016), respectively. The M57T substitution is highlighted in orange.

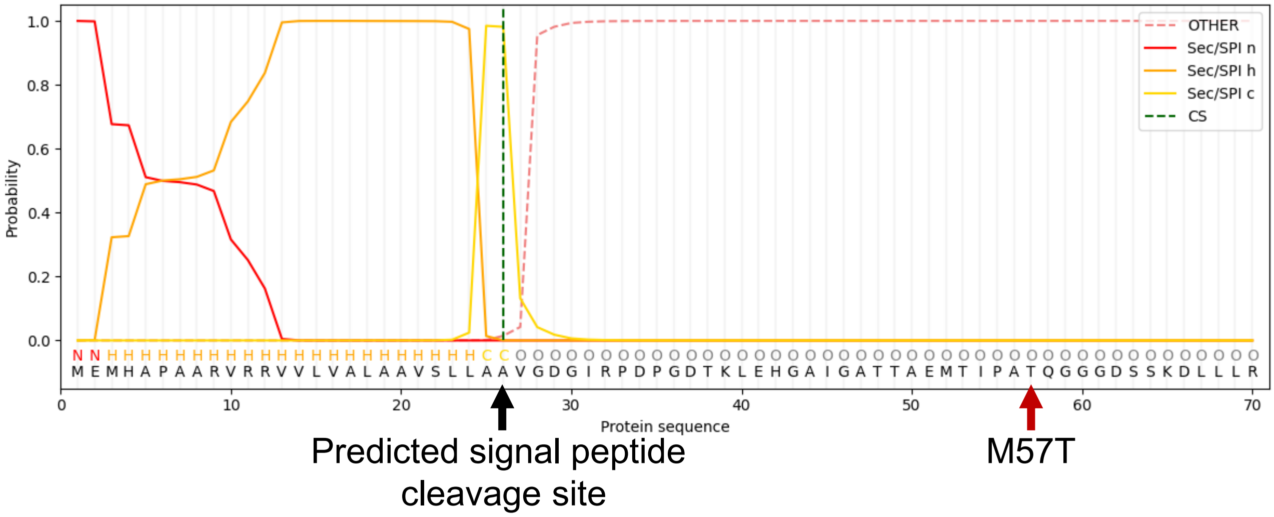


**Supplementary Fig. S21 Prediction of the signal peptide in PgEPF2**
Signal peptide prediction for PgEPF2 using SignalP. The predicted signal peptide cleavage site is indicated by a black arrow. The position of the M57T substitution identified in PgEPF2 is also indicated. The x-axis represents amino acid position, and the y-axis represents the probability assigned to each SignalP prediction class.
